# Hapln1-HA signaling promotes progenitor cell proliferation and spinal cord regeneration

**DOI:** 10.64898/2026.03.28.714985

**Authors:** Yuxiao Xu, Lili Zhou, Vishnu Saraswathy, Ryan McAdow, Mayssa H. Mokalled

## Abstract

Adult zebrafish exhibit scarless repair and functional recovery following spinal cord injury. Their regenerative capacity is attributed to potent stem-like progenitors that mediate neuronal and glial repair. Zebrafish are thought to lack anti-regenerative extracellular matrix (ECM) components abundant in mammalian SCI, but the positive contributions of ECM to spontaneous spinal cord repair are less understood. By employing cross-species single-cell transcriptomics, we found the hyaluran modifying enzyme Hapln1 is upregulated in zebrafish progenitors but not in mouse progenitors following injury. Loss-of-function of *hapln1a/b* and ablation of *hapln1*^+^ cells reduce progenitor cell activation and hinder spontaneous recovery from injury. Using a series of *in vivo* and *in vitro* assays, we show that Hapln1 is required for hyaluran-*cd44b* mediated progenitor cell proliferation. This study reveals that, in addition to lacking anti-regenerative ECM components around SC lesions, zebrafish can also leverage pro-regenerative ECM molecules to enhance progenitor cell potency and promote repair.

## INTRODUCTION

Regenerative capacity is unevenly distributed among vertebrates. In most mammalian species including humans, spinal cord injury (SCI) causes permanent functional impairment and long-term burdens (Silver, 2016; Sofroniew, 2018; Zheng and Tuszynski, 2023). Conversely, adult zebrafish undergo spontaneous repair and recover swim function following SCI (Becker and Becker, 2008; Tendolkar and Mokalled, 2025). The presence of robust progenitor cells and the lack of scarring distinguish zebrafish from mammalian SCI and underlie their spontaneous repair following injury.

Spinal cord (SC) repair in adult zebrafish is enabled by potent stem cell-like progenitors (Hui et al., 2010; Weinholtz et al., 2026). Zebrafish SC progenitors express stem cell markers like *sox2* and were recently found to comprise ependymal and parenchymal subtypes (Weinholtz et al., 2026). *sox2*^+^ progenitors are compartmentalized into lineage-restricted niches that transiently proliferate and differentiate into neurons and glia following injury (Reimer et al., 2008; Mokalled et al., 2016; Klatt Shaw et al., 2021; Saraswathy et al., 2022; Zhou et al., 2023). In contrast, non-regenerative mammalian SCs rely on adult ependymal cells that proliferate but primarily differentiate into astrocytes and oligodendrocytes after injury (Horky et al., 2006; Meletis et al., 2008; Barnabe-Heider et al., 2010; Sabelström et al., 2013). We propose that elucidating transcriptional similarities and differences between zebrafish and mammalian progenitors will uncover molecular programs that underlie heightened stem cell potency in zebrafish, which will inform and advance stem cell-based interventions for mammalian SCI.

Often studied in the context of anti-regenerative scarring, the extracellular matrix (ECM) undergoes dynamic changes following SCI. Consistent with their inhibitory roles in the lesion environment, the majority of ECM molecules studied in SCI are typically upregulated in mammalian SC lesions and impede SC repair (Lemons et al., 1999; Jones et al., 2003; Kolb et al., 2023). Comparative proteomics between rodents and zebrafish identified inhibitory factors that occupy mammalian but not zebrafish SC lesions (Kolb et al., 2023). Exogenous expression of these factors in zebrafish is anti-regenerative, suggesting the absence of inhibitory ECM underlies spontaneous SC repair in zebrafish (Kolb et al., 2023). Recent zebrafish studies support a complementary model in which ECM molecules such as chondroitin sulfate proteoglycans (CSPGs) are anti-regenerative in mammals but possess pro-regenerative functions in zebrafish (Kisucká et al., 2026; Lafouasse et al., 2026). CSPGs accumulate after mammalian SCI where they impede SC repair. Intriguingly, enzymatic removal of CSPGs is sufficient to alleviate injury symptoms in mammals but was recently reported to show anti-regenerative effects zebrafish (Kisucká et al., 2026; Lafouasse et al., 2026). While prior research has uncovered anti-regenerative roles for the ECM within SC lesions, whether and how zebrafish leverage ECM molecules to promote progenitor cell potency and SC repair is less understood.

Hyaluronan and proteoglycan link protein 1 (Hapln1) is a crucial ECM organizing factor that stabilizes proteoglycan-hyaluronan complexes (Long et al., 2018). Hyaluronan, also known as hyaluronic acid (HA), is a glycosaminoglycan that controls ECM stiffness and regulates various biological processes. Native HA is a high molecular weight polymer that activates Cd44-mediated signaling (Cyphert et al., 2015). HA binds Cd44 receptor complexes, activating PI3K or ERK signaling and directing cell survival, proliferation or differentiation in a context-dependent manner. In the mammalian CNS, HA is primarily studied in the contexts of astrocyte proliferation and neural net integrity during glioblastoma or following injury (Wolf et al., 2020; Grycz et al., 2022; Sanchez-Ventura et al., 2022). Mechanistic insights into the roles of Hapln1/HA in SC progenitors could be inferred from brain development, whereby HA is abundantly present around neurogenic stem cells during embryonic development and is postnatally decreased in mice (Margolis et al., 1975; Preston and Sherman, 2011). *hapln1* genes have been implicated in cardiac regeneration and vascular growth in zebrafish, but their roles in SC tissues or SCI are less understood (Mahony et al., 2021; Sun et al., 2022; Liu et al., 2023; Sun et al., 2023).

Here, cross-species single-cell comparisons show zebrafish spinal progenitors exhibit greater similarity to neurogenic progenitors in the mouse brain than to mouse spinal progenitors. Following injury, SC progenitors upregulate ECM-related genes including *hapln1* in zebrafish but not in mice. Loss-of-function of *hapln1a/b* and ablation of *hapln1*^+^ cells show *hapln1-cd44b* signaling is required for proliferative activation of *cd44b*^+^ progenitors and spontaneous repair following injury. *in vivo* and *in vitro* assays establish a requirement for Hapln1 in HA-mediated signaling and progenitor cell proliferation. Thus, not only do zebrafish lack anti-regenerative ECM within SC lesions, but they also harness pro-regenerative ECM to support progenitor cell potency and tissue repair.

## RESULTS

### Zebrafish and mouse SC progenitors possess distinct molecular profiles

We compared *sox2*^+^ progenitors from zebrafish and mice to identify molecular differences that may contribute to their differential regenerative capacity (**Fig. 1A**). Single-cell RNA-sequencing datasets comprising adult SC tissues from zebrafish and mice after SCI were used (Milich et al., 2021; Weinholtz et al., 2026). As the mammalian brain is more permissive for adult neurogenesis than the mammalian SC, αSMA-labelled *sox2*^+^ ependymal cells and αSMA^-^ *sox2*^+^ progenitors from the subventricular zone of uninjured mouse brains were included as controls (Shah et al., 2018). In a first analysis, we subset and integrated *sox2*^+^ cells from mouse SCI at 0, 1, 3 and 7 dpi with *sox2*^+^ cells from zebrafish SCI at 0, 7 and 14 dpi. This analysis yielded 28 clusters (**Fig. 1B, S1A)**. We found *sox2*^+^ cells from mouse and zebrafish SCs possessed distinct transcriptional signatures, regardless of injury time point (**Fig. 1B**). Clusters where either zebrafish or mouse cells comprised <25% of total cell numbers were classified as species-specific. On the hand, clusters where zebrafish and mouse cells comprised >25% of total cell clusters were classified as “shared”. Out of 28 clusters, only clusters 5, 11 and 12 exhibited transcriptional identities that were shared between zebrafish and mice (**Fig. 1C, S1B)**. Among the remaining 25 species-specific clusters, 7 clusters were classified as zebrafish-specific and 18 clusters were mouse-specific (**Fig. 1C**). These cross-species differences were even more pronounced when comparing matched time points (**Fig. 1D, 1E)**. For instance, *sox2*^+^ cells from either uninjured or 7 dpi SC tissues showed little transcriptional similarity between zebrafish and mice (**Fig. 1D, 1E)**. Notably, the molecular profiles of murine *sox2*^+^ cells exhibited drastic transcriptional changes from 0 to 7 dpi (**Fig. 1D, 1E)**. However, compared to mouse cells, changes to the molecular identities of zebrafish *sox2*^+^ cells were less pronounced at 7 dpi compared to uninjured controls (**Fig. 1D, 1E)**. These results are consistent with recent observations that *sox2*^+^ cells are heterogeneous and lineage-biased before and after SCI (Weinholtz et al., 2026), suggesting *sox2*^+^ cells may be poised to respond to injury and support repair with little transcriptional changes in zebrafish. Overall, these findings revealed differences in progenitor cell identities between innately regenerative zebrafish and poorly regenerative mice.

**Figure 1.**
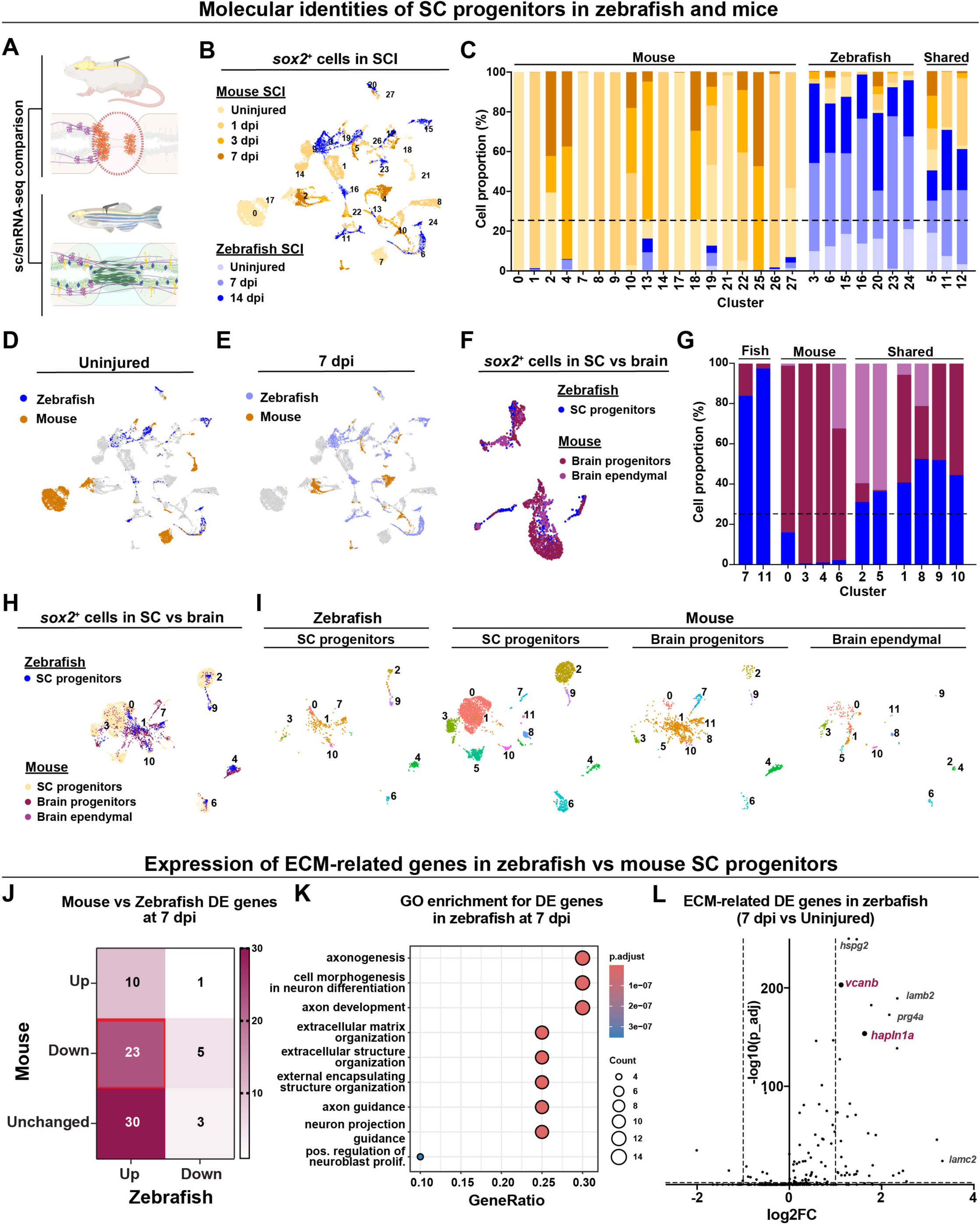
Comparative transcriptomics between zebrafish and mouse SC progenitors. **(A)** snRNA-seq datasets for zebrafish and mouse SCI were used to compare *sox2*^+^ cells from regenerative and non-regenerative species. **(B-E)** Integration of *sox2*^+^ cells from zebrafish and mouse SCs following SCI. UMAP of the integrated dataset highlighting all cells (B), uninjured cells (D) or cells at 7 dpi (E). Clusters were classified as mouse-only, zebrafish-only or shared (C). **(F, G)** UMAP of the integrated dataset of SC progenitors from uninjured zebrafish with progenitor or ependymal cells from the mouse brain (F). Clusters were classified as mouse-only, zebrafish-only or shared (G). **(H, I)** Combined (H) and split (I) UMAP of the integrated dataset of SC progenitors from uninjured zebrafish, SC progenitors from uninjured mice, with progenitor or ependymal cells from the mouse brain. **(J)** Heatmap shows ECM markers with log_2_(FC)>1 in zebrafish SCs at 7 dpi. Grid color represents the number of markers in the corresponding category. **(K)** Dot plot of GO terms enriched in *cd44b*^+^ *sox2*^+^ cells. Dot size and color represent percent expressing cells and average gene expression, respectively. **(L)** Volcano plot of differentially expressed ECM-related genes at 7 dpi versus uninjured controls in zebrafish SCs.

We postulated the disparities in molecular identities between zebrafish and mouse SC progenitors could be attributed to species-driven molecular differences or divergent SCI responses. To address these possibilities, we integrated *sox2*^+^ cells from uninjured zebrafish SCs with progenitor and ependymal cells from uninjured mouse brain (**Fig. 1F, S1C)** (Shah et al., 2018). Intriguingly, out of 12 clusters identified in this integration, 6 clusters (cluster 1, 2, 5, 8, 9 and 10) were shared between zebrafish and mouse cells (**Fig. 1G, S1D)**. Finally, we performed a 4-way integration to compare *sox2*^+^ cells from uninjured zebrafish SCs to *sox2*^+^ cells from uninjured mouse SCs, in addition to progenitor and ependymal cells from uninjured mouse brain (**Fig. 1H**). This integration confirmed zebrafish SC progenitors are more similar to neurogenic mouse brain progenitors than to mouse SC progenitors (**Fig. 1H**). We conclude that species or technical differences are unlikely to underlie the different identities observed between zebrafish and mouse progenitors, and that the transcriptional similarities observed in our cross-species integrations correlate with the neurogenic potential of the progenitor populations included in this analysis.

We analyzed the gene ontology (GO) for the top markers from each cluster in the integration of zebrafish and mouse SC progenitors and observed 2 major species-driven clusters **(Fig. S1E)**. One set of GO terms were preferentially expressed in mouse SC progenitors and related to metabolism and cell synthesis (e.g. metabolism of RNA, cell cycle and ribosome biogenesis) **(Fig. S1E)**. The second set of GO terms were identified in zebrafish or shared SC progenitors and were associated with neuronal function (e.g. neuron projection morphogenesis, synapse organization and modulation of chemical synaptic transmission) **(Fig. S1E)**. This analysis underscored the neurogenic potential and molecular identities associated with spontaneous tissue repair in zebrafish progenitors compared to mice. Together, these studies showed zebrafish and mouse SC progenitors possess intrinsic transcriptional differences at baseline and after SCI and suggested these cell identity differences may underlie differential stem cell potency and regenerative capacity across species.

### Zebrafish SC progenitors upregulate ECM-related genes after SCI

Previous proteomic studies reported that the majority of lesion-associated ECM components are upregulated in rat SCI and downregulated in zebrafish SCI (Kolb et al., 2023). These observations are consistent with ECM being anti-regenerative, hence downregulated in regenerative zebrafish. However, noting that multiple ECM-related genes were expressed in the transcriptomes of zebrafish SC progenitors, we postulated zebrafish progenitors may leverage pro-regenerative ECM to direct spontaneous SC repair. To achieve in depth understanding of ECM functions in SC progenitors, we compiled a comprehensive list of zebrafish and mouse ECM-related genes from Matrisome.org **(Table S1)** (Naba et al., 2012; Nauroy et al., 2018). Pseudobulk analysis of the integrated dataset of zebrafish and mouse SC progenitors identified 63 ECM genes that were upregulated in zebrafish SC progenitors at 7 dpi compared to uninjured controls **(Table S2)**. Among these 63 genes, 23 were downregulated and 30 were not significantly changed in mice at 7 dpi (**Fig. 1J**). Intriguingly, only 10 out of 63 ECM genes that were upregulated in zebrafish progenitors were also upregulated in mouse progenitors at 7 dpi (**Fig. 1J**). On the other hand, 9 ECM-related genes were downregulated in zebrafish progenitors, only 1 of which was upregulated in mouse progenitors (**Fig. 1J**). Thus, unlike the zebrafish lesion environment where inhibitory ECM molecules are not expressed, zebrafish SC progenitors upregulate ECM-related genes that are downregulated or absent in zebrafish progenitors following SCI.

Gene ontology of the 53 genes that were upregulated in zebrafish at 7 dpi but downregulated or unchanged in mice revealed terms related to neural development (axonogenesis, axon development and neuron projection guidance) and stem cell function (positive regulation of neuroblast proliferation), in addition to terms related to general ECM functions (extracellular structure organization) (**Fig. 1K**). Genes upregulated following SCI included *col2a1a, hapln1a, hspg2, igfbp3, lamb2, lamc2, prg4at, rspo2* and *vcanb* (**Fig. 1L**). Main motifs emerging from this analysis were genes encoding for HA-binding proteins (*hapln1a, prg4a*, *bcan* and *vcanb*) and laminins (*lamb2, lamb4* and *lamc2*). These gene expression changes were concomitant with acute progenitor cell proliferation in the first 7 days following SCI, and are therefore likely to play pro-regenerative roles during spontaneous SC repair.

### *hapln1a* is upregulated in zebrafish progenitors following SCI

Genes encoding Hapln1 (*hapln1* in mice and *hapln1a*/*b* in zebrafish) were differentially regulated in zebrafish and mouse SCI. In RNA-seq datasets encompassing all spinal cell types, murine *hapln1* expression was negligible and unchanged across time points, whereas zebrafish *hapln1a* and *b* were upregulated at 7 and 14 dpi relative to controls (**Fig. 2A**). We next investigated *hapln1* expression in our integrated dataset of zebrafish and mouse SC progenitors. Noting that an essential step in this cross-species integration is the conversion of zebrafish genes into their murine orthologs, both zebrafish *hapln1a* and *b* were represented by a single *hapln1* ortholog in this analysis. These gene expression studies confirmed that *hapln1* is preferentially upregulated in zebrafish SC progenitors but not in mouse SC progenitors after injury (**Fig. 2B**). Intriguingly, in our integrated dataset of uninjured SC progenitors from zebrafish with murine brain cells, *hapln1* was preferentially expressed in clusters shared between zebrafish cells and neurogenic brain progenitors (clusters 1, 8, 9, 10), but not in clusters shared between zebrafish SCs and non-neurogenic brain ependymal cells (clusters 2, 5) (**Fig. 2C**). These findings showed *hapln1* is upregulated in spinal progenitors after zebrafish SCI and suggested *hapln1* regulation may correlate with neurogenic stem cell potential in zebrafish SCs.

**Figure 2.**
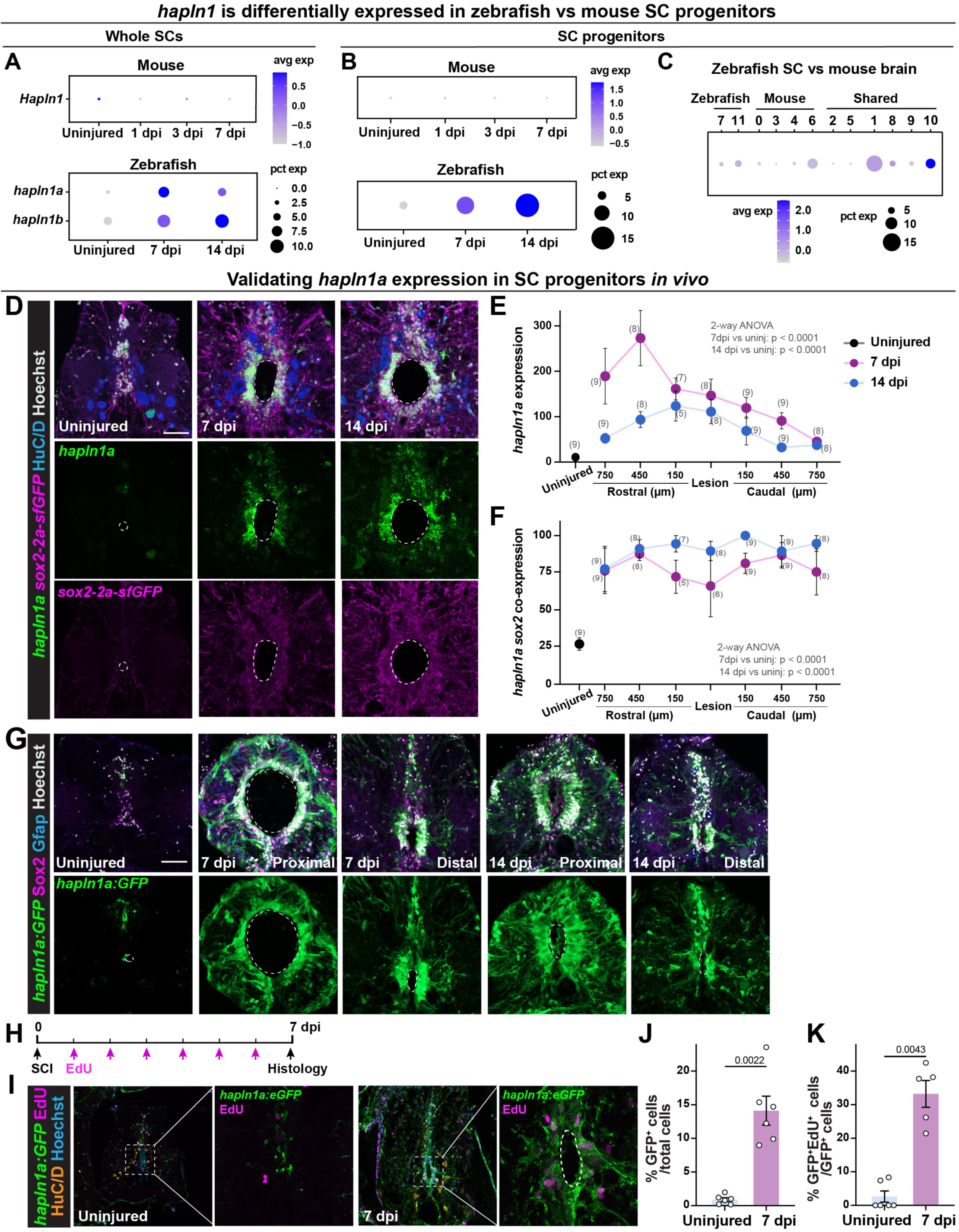
*hapln1* is upregulated in progenitor cells following SCI. **(A)** Dot plot shows *hapln1* expression in whole SCs following zebrafish or mice SCI. **(B)** Dot plot shows *hapln1* expression in *sox2*^+^ cells from zebrafish or mouse SCs. **(C)** Dot plot shows *hapln1* expression in the integrated dataset of SC progenitors from uninjured zebrafish with progenitor or ependymal cells from the mouse brain. For all dot plots, dot size and color represent percent expressing cells and average gene expression, respectively. **(D)** HCR *in situ* hybridization for *hapln1a* in *sox2-2a-sfGFP* zebrafish at 0, 7 and 14 dpi. SC cross-sections 450 µm rostral to the lesion are shown. **(E, F)** Quantification of *hapln1a* expression (E) and hapln1 co-localization with *sox2-2a-sfGFP* (F). Multiple tissue levels 750 µm rostral and caudal to the the lesion were quantified. **(G)** hapln1a-driven expression in *hapln1a:GFP* reporter zebrafish. SC cross sections 450 µm (proximal) and 750 µm (distal) from the lesion are shown at 7 and 14 dpi. Sections were stained for Sox2 and Gfap. **(H-K)** Experimental timeline to assess EdU incorporation in *hapln1a:GFP* reporter fish (H). GFP and EdU staining was performed (I). The proportions of GFP^+^ cells (J) and GFP^+^ EdU^+^ cells within GFP^+^ cells (K) were quantified. For all micrographs, dashed lines delineate the central canal. Data points represent individual animals and n numbers are indicated in parentheses. Statistical tests and p-values are indicated. Scale bars: 50 µm.

We performed Hybridization Chain Reaction (HCR) *in situ* hybridization for *hapln1a* and *b* to examine their expression in regenerating SC tissues. To this end, *sox2-2a-sfGFP* knock-in zebrafish were subjected to complete SC transections, and *hapln1* transcripts were co-stained with GFP and HuC/D to respectively label *sox2*^+^ progenitors and HuC/D^+^ neurons (**Fig. 2D**) (Shin et al., 2014). Consistent with bioinformatics data (**Fig. 2A, 2B)**, *hapln1a* expression was negligible in uninjured SCs and was markedly upregulated at 7 and 14 dpi (**Fig. 2D**). *hapln1a* transcripts were predominant in lateral ependymal progenitors around the central canal, with low transcript levels in dorsoventral progenitors and SC parenchyma (**Fig. 2D**). Quantification showed *hapln1a* expression peaked at 7 dpi and co-localized with *sox2* following (**Fig. 2E, 2F)**. Compared to *hapln1a* (**Fig. 2D**)*, hapln1b* transcripts were less expressed and predominantly detected in parenchymal *sox2*^+^ cells rather than ependymal cells **(Fig. S2A)**. These results established *hapln1* genes are upregulated in *sox2*^+^ cells after SCI.

### *hapln1a^+^* progenitors are proliferative after SCI

We used an established *hapln1a:GFP* transgenic zebrafish line to characterize the dynamics of *hapln1a*-driven expression during SC regeneration (Sun et al., 2022). Validating our *in situ* hybridization, SC cross sections from *hapln1a:GFP* showed GFP was strongly induced after SCI (**Fig. 2G**). To examine the stem cell properties of *hapln1a*^+^ cells during SC regeneration, adult *hapln1a:GFP* zebrafish were subjected to complete SC transections followed by daily intraperitoneal EdU injections after injury (**Fig. 2H**). This EdU labeling regimen establishes a cumulative profile of injury-induced cell proliferation. SC tissues at 7 dpi were compared to tissues collected from uninjured siblings after 6 days of daily EdU injection (**Fig. 2I**). SC cross sections 450 µm rostral to the lesion were analyzed. Compared to uninjured controls where 0.89% of SC cells were GFP^+^, *hapln1a:GFP* robustly increased and accounted for 14.1% of total cells at 7 dpi (**Fig. 2J, S2B)**. Mirroring global cell proliferation, the proportion of GFP^+^ EdU^+^ cells increased from 2.62% in uninjured controls to 33.19% at 7 dpi (**Fig. 2K, S2C-E)**. These findings established *hapln1a*^+^ cells are relatively quiescent in homeostatic adult SCs and mount an acute proliferative response within 7 days of SCI.

### *hapln1a*^+^ cells are required for SC regeneration

To investigate if *hapln1a*^+^ progenitors are required for regeneration, we used *hapln1a*:mCherry-Nitroreductase (*hapln1a*:mCherry-NTR) transgenic animals for cell ablation studies (Sun et al., 2022). In this system, *hapln1a*-driven expression of the bacterial NTR enzyme was used to catalyze the reduction of the prodrug Metrodinazole (MTZ) into a cytotoxic product that induces cell death (Curado et al., 2008). To ablate *hapln1a*^+^ progenitors, we subjected *hapln1a*:mCherry-NTR (Tg^+^) fish to complete SC transections, followed by 3 consecutive treatments with 5 mM MTZ MTZ for 14 hrs at 5, 6 and 7 dpi (**Fig. 3A**). Immunohistochemistry for cleaved caspase 3 (Ccp3) was performed at 8 dpi to assess the extent of cell death using this ablation regimen (**Fig. 3B**). Concomitant with increased cell death, mCherry expression was fragmented and markedly reduced in MTZ-treated NTR fish (**Fig. 3B**). The profiles of Ccp3^+^ cells were increased in MTZ-treated NTR animals compared to vehicle-treated siblings (**Fig. 3C**). These studies established a genetic ablation model for *hapln1a*^+^ cells and were subsequently used to examine their requirement during SC regeneration.

**Figure 3.**
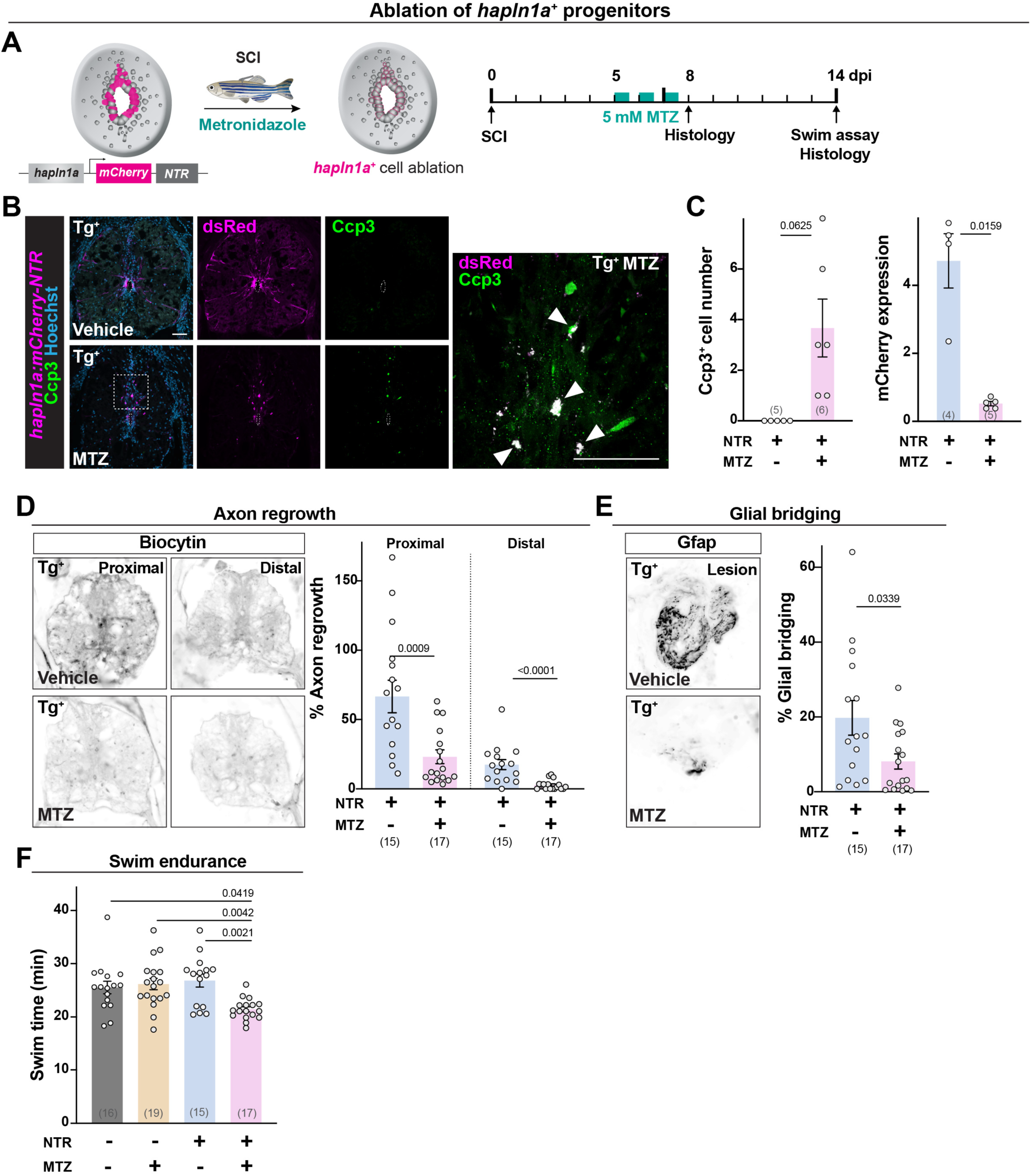
Ablation of *hapln1a*^+^ cells impairs SC repair. **(A)** Experimental timeline to induce *hapln1a*^+^ cell ablation *hapln1a:mCherry-NTR* zebrafish. **(B, C)** mCherry expression and staining for cleaved Caspace 3 (Ccp3) were performed at 8 dpi. **(D)** Anterograde axon tracing in MTZ- or vehicle-treated *hapln1a:mCherry-NTR* (Tg^+^) fish at 14 dpi. Representative images show the extent of Biocytin caudal to the lesion. Axon growth was first normalized to the extent of Biocytin rostral to the lesion, and then to labeling in controls. **(E)** Glial bridging following *hapln1a*^+^ cell ablation at 14 dpi. Representative images show the Gfap^+^ bridge at the lesion site. Percent bridging represents the cross-sectional area of the glial bridge at the lesion site relative to the intact SC 750 µm rostral to the lesion. **(F)** Swim endurance following *hapln1a*^+^ cell ablation at 14 dpi. MTZ-treated Tg^+^ animals are shown in addition to vehicle and Tg^-^ control groups. This behavioral test assessed the capacity of animals to swim in an enclosed swim tunnel under increasing water current velocities. Time at exhaustion is shown. Data points represent individual animals and n numbers are indicated in parentheses. Statistical tests and p-values are indicated.

To determine the impact of *hapln1a*^+^ cell ablation during SC regeneration, we subjected *hapln1a*:mCherry-NTR (Tg^+^) fish to complete SC transections and MTZ treatment. Glial bridging, axon regrowth and swim endurance were then performed at 14 dpi. For anterograde axon tracing, Biocytin was applied at the hindbrain level and Biocytin-labeled axons were traced at 600 μm (proximal) and 1500 μm (distal) caudal to the lesion (**Fig. 3D**). In this assay, axon regrowth was 2.4-fold attenuated in proximal SC tissues from MTZ-treated Tg^+^ animals compared to MTZ-treated Tg^-^ controls (**Fig. 3D**). For glial bridging, Gfap immunostaining was used to determine the areas of glial bridges at the lesion site relative to the areas of intact SC tissues (**Fig. 3E**). Percent bridging at the lesion site decreased 2.8-fold in MTZ-treated Tg^+^ animals compared to MTZ-treated Tg^-^ controls (**Fig. 3E**). To evaluate the functional impact of ablating *hapln1a*^+^ cells after SCI, we assessed the swim capacities of ablated animals in an enclosed swim tunnel under increasing water current velocities (Burris et al., 2021; Burris and Mokalled, 2024). Swim endurance was significantly diminished in MTZ-treated Tg^+^ animals compared to MTZ-treated Tg^-^and vehicle-treated Tg^+^ controls (**Fig. 3F**). These experiments established an essential role *hapln1a*^+^ cells during SC regeneration in zebrafish.

### *hapln1a/b* mutants show impaired SC regeneration

To examine the role of *hapln1a*/b during functional recovery, we performed SC transections on *hapln1a* and *b* double mutant zebrafish, referred to hereafter as *hapln1a/b* mutants or *hapln1a/b^-/-^* (Sun et al., 2022). We then evaluated functional recovery at 14, 28 and 42 dpi (**Fig. 4A**). Double wild-type siblings for *hapln1a* and *b* (*hapln1a/b^+/+^*) were used as controls. Swim endurance was comparable across cohorts of uninjured fish and declined in both groups after SCI. However, *hapln1a/b* mutants showed significantly lower swim endurance compared to controls at 42 dpi (**Fig. 4A**). Swim endurance was unaffected in single homozygous mutants for either *hapln1a* or *hapln1b* compared to their control siblings **(Fig. S3A)**, likely due to genetic compensation among gene paralogs. To assess anatomical regeneration in *hapln1a/b* mutants, rostral neurons were anterogradely traced with biocytin and visualized caudal to the lesion. At 42 dpi, the proportion of caudally traced axons proximal to the lesion and glial bridging were significantly reduced in *hapln1a/b^-/-^*compared to wild-type siblings (**Fig. 4B, 4C)**. These results showed *hapln1a/b* are required for functional recovery and anatomical regeneration following SCI.

**Figure 4.**
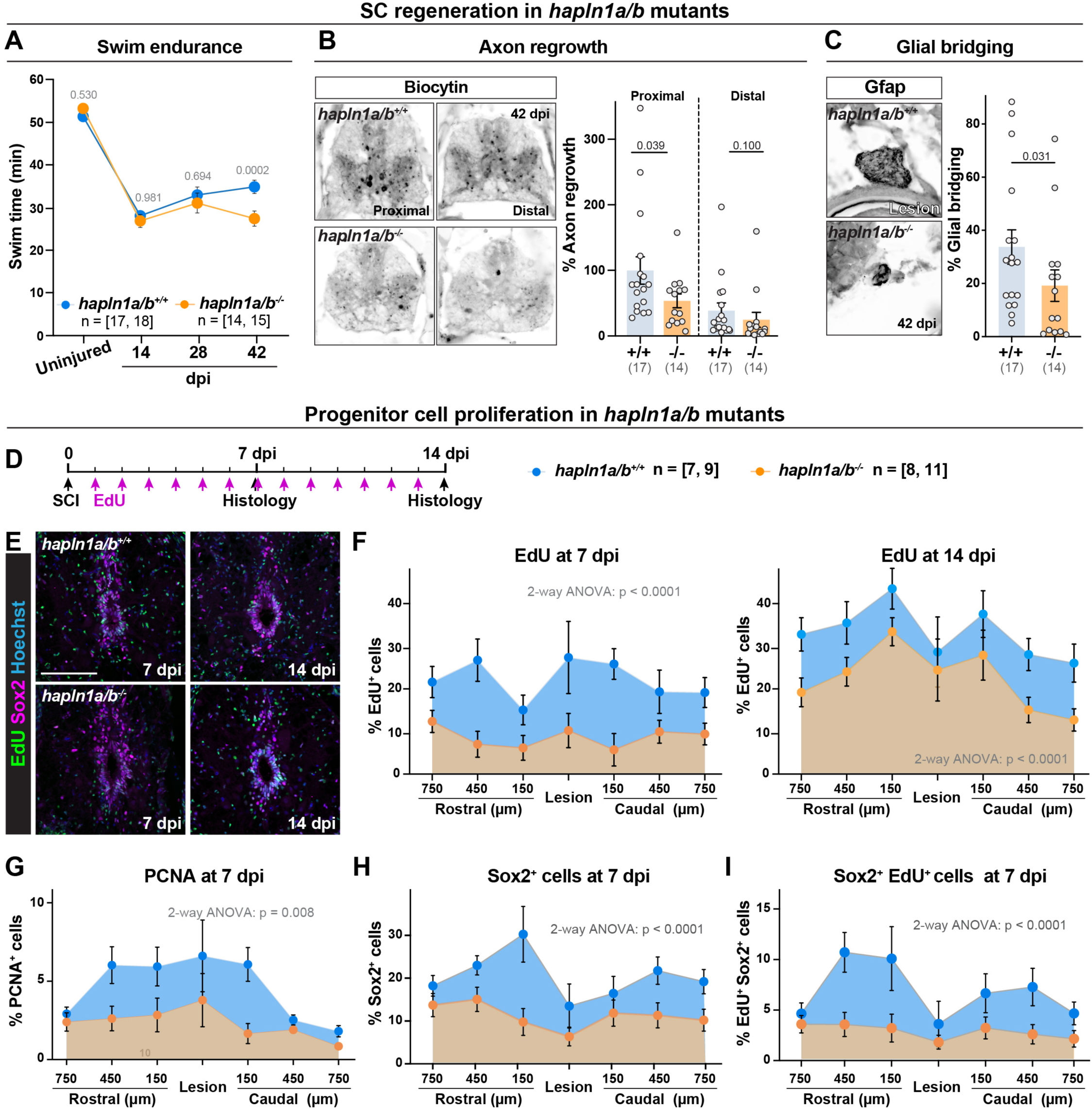
*hapln1a/b* exhibit impaired progenitor cell proliferation and SC regeneration. **(A)** Swim endurance in *hapln1a/b*^-/-^ and wild-type siblings at 0, 14, 28 and 42 dpi. **(B)** Anterograde axon tracing in *hapln1a/b*^-/-^ and wild-type siblings at 42 dpi. Anterograde biocytin tracer was administered to each fish at 42 dpi. Representative images show the extent of Biocytin caudal to the lesion. Axon growth was first normalized to the extent of Biocytin rostral to the lesion, and then to labeling in controls. **(C)** Glial bridging *hapln1a/b*^-/-^ and wild-type siblings at 42 dpi. Representative images show the Gfap^+^ bridge at the lesion site. Percent bridging represents the cross-sectional area of the glial bridge at the lesion site relative to the intact SC 750 µm rostral to the lesion. **(D)** Experimental timeline to assess EdU incorporation in *hapln1a/b* mutants. **(E)** Sox2 and EdU staining was performed at 7 dpi and 14 dpi. SC cross sections 450 µm from the lesion are shown. **(F-I)** Quantification of EdU incorporation (F), PCNA^+^ cells (G), Sox2^+^ cells (H) and Sox2^+^ EdU^+^ cells (I) at 7 or 14 dpi. Multiple tissue levels 750 µm rostral and caudal to the lesion were quantified. Data points represent individual animals in B and C and n numbers are indicated. Statistical tests and p-values are indicated. Scale bars: 100 µm.

### *hapln1a/b* loss-of-function impairs progenitor activation and neurogenesis

As *hapln1* expression is upregulated in *sox2*^+^ progenitors after SCI (**Fig. 2**), we postulated that defective progenitor cell activation may underlie *hapln1a/b* mutant phenotypes. To examine the role of *hapln1* in progenitor activation, we assayed the proliferation of Sox2^+^ cells in *hapln1a/b* mutants. To this end, *hapln1a/b^-/-^*and wild-type siblings were subjected to SCI and daily EdU injections (**Fig. 4D**). Sox2 and EdU staining were performed at 7 and 14 dpi (**Fig. 4E**). SC sections 750, 450 and 150 µm rostral and caudal to the lesion were analyzed. This analysis showed EdU incorporation was significantly decreased in *hapln1a/b* mutants at 7 and 14 dpi (**Fig. 4F, S3)**. Noting that defects in cell proliferation were more prominent at 7 dpi compared to 14 dpi, we performed PCNA staining to assess snapshots of actively dividing cells at 7 and 14 dpi. The numbers and proportions of PCNA^+^ cells were significantly decreased in *hapln1a/b* mutants at 7 dpi but not at 14 dpi (**Fig. 4G, S3C, S3D**). Consistent with decreased progenitor cell proliferation, the proportions of Sox2^+^ and EdU^+^ Sox2^+^ cells were reduced in *hapln1a/b* mutants at 7 dpi and unchanged at 14 dpi (**Fig. 4H, S3E, S3F**). These findings showed *hapln1* is required for acute proliferative activation of *sox2*^+^ progenitors during the first 7 days following SCI.

### *hapln1a/b* loss-of-function impairs the proliferation of *cd44b*^+^ progenitors

To elucidate the mechanism by which Hapln1 impacts progenitor cell proliferation, we examined the expression of the HA receptor Cd44 (**Fig. 5A**). By *in situ* hybridization, we observed baseline expression of *cd44b* transcripts in uninjured control SCs (**Fig. 5B**). Concomitant with progenitor cell proliferation following SCI, *cd44b* expression was markedly upregulated in *sox2-2a-sfGFP* cells at 7 and 14 dpi (**Fig. 5B, 5C)**. To evaluate the anatomical organization of *cd44b^+^ sox2^+^*progenitors relative to *hapln1* expression, we subjected *hapln1a:GFP* transgenic fish to SC transections, collected SC tissues at 7 dpi and performed *in situ* hybridization for *cd44b* (**Fig. 5D**). This experiment revealed 2 patterns of *cd44b* expression: weak expression in *hapln1a:GFP* cells lateral to the central canal and strong expression in cells adjacent to *hapln1a:GFP* cells dorso-lateral and ventro-lateral to the central canal (**Fig. 5D**). Our snRNA-seq data showed *cd44b* expression in SC tissues at baseline and following SCI, whereas its paralog *cd44a* showed markedly lower expression before and after injury (**Fig. 5E**). Supporting a model where *cd44b^+^* progenitors are poised to respond to injury in homeostatic SCs*, cd44b* was expressed in similar proportions of *sox2^+^*progenitors across uninjured, 7 and 14 dpi tissues (**Fig. 5E**). *cd44b^+^ sox2^+^* cells expressed genes related to axonogenesis, axon development, axon guidance and SC development (**Fig. 5F).** The proximity of *hapln1* and *cd44b* expressing cells supported a potential role for Hapln1-HA-Cd44b signaling in progenitor cell activation following SCI.

**Figure 5.**
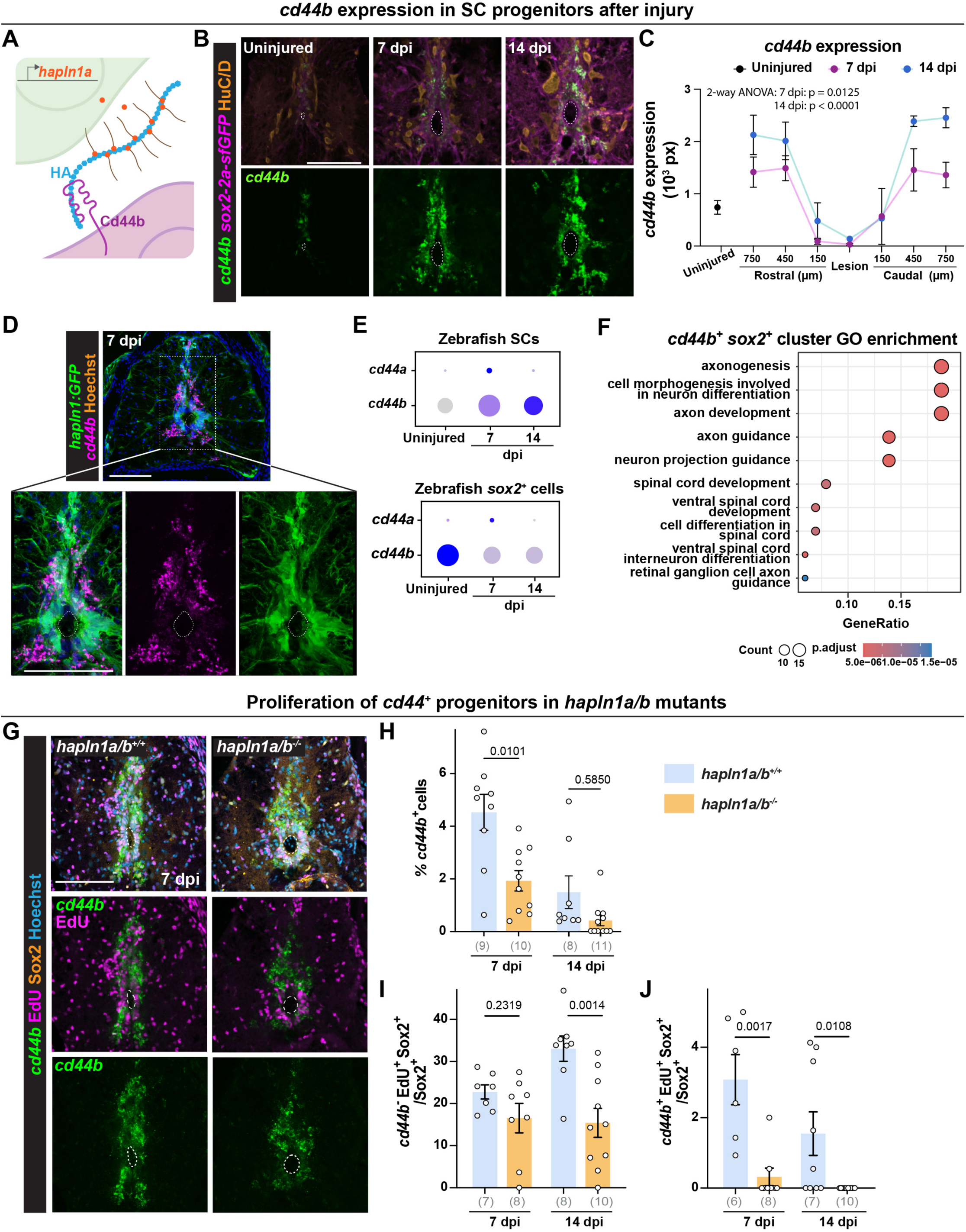
*hapln1a/b* exhibit altered HA deposition and the proliferation of *cd44b^+^* progenitors. **(A)** Our working model is that Hapln1 stabilizes HA and HA-dependent signaling via Cd44 receptor complexes. **(B, C)** HCR *in situ* hybridization for *cd44b* in *sox2-2a-sfGFP* fish at 0, 7 and 14 dpi. SC cross sections 450 µm from the lesion are shown. Multiple tissue levels 750 µm rostral and caudal to the lesion were quantified. **(D)** HCR *in situ* hybridization for *cd44b* in *hapln1a:GFP* fish at 7 dpi. **(E)** Dot plot shows cd44a and b expression in whole zebrafish SCs (upper) or in subsets of *sox2*^+^ cells (lower). Dot size and color represent percent expressing cells and average gene expression, respectively. **(F)** Gene Ontology of genes enriched in *cd44b^+^ sox2^+^* cells. **(G)** HCR *in situ* hybridization for *cd44b* and staining for EdU and Sox2 were performed. *hapln1a/b*^-/-^ and wild-type siblings were subjected to daily EdU injections and analyzed at 7 dpi. **(H-J)** The proportions of *cd44b*^+^ cells (H), *cd44b*^-^ EdU^+^ Sox2^+^ cells relative to Sox2^+^ cells (I) and *cd44b*^+^ EdU^+^ Sox2^+^ cells relative to Sox2^+^ cells (H) were quantified. For all micrographs, dashed lines delineate the central canal. Data points represent individual animals and n numbers are indicated in parentheses. Statistical tests and p-values are indicated. Scale bars: 50 µm.

To evaluate how *hapln1a/b* deletion impacts the proliferation of *cd44b*^+^ cells, we performed SC transections and daily EdU injections on *hapln1a/b^-/-^* and wild-type controls. SC tissues were collected at 7 and 14 dpi, and *cd44b in situ* hybridization was combined with Sox2 and EdU staining (**Fig. 5G**). Recapitulating our prior observation, EdU incorporation and sox2+ cells were decreased in *hapln1a/b^-/-^*fish **(Fig. S4A-D)**. The numbers and proportions of *cd44b*^+^ cells were significantly reduced in *hapln1a/b* mutants at 7 dpi (**Fig. 5H, S4E)**. Compared to 7 dpi, *cd44b*^+^ cells decreased in wild-type at 14 dpi, suggesting *cd44b*^+^ progenitors are early responders that are activated in the first 7 days post-injury (**Fig. 5H, S4E)**. A less pronounced reduction in *cd44b*^+^ cells was observed in *hapln1a/b* mutants compared to wild types at 14 dpi (**Fig. 5H, S4E)**. To evaluate whether *hapln1* loss affects the proliferation of *cd44b^+^* progenitors, we quantified EdU incorporation in *cd44b*^-^ *sox2*^+^ and in *cd44b*^+^ *sox2*^+^ cells. This analysis revealed cell proliferation is acutely reduced in *cd44b*^+^ *sox2*^+^ cells but not in *cd44b*^-^ *sox2*^+^ at 7 dpi (**Fig. 5I, 5J S4F, S4G)**. These results indicated *hapln1a/b* are required for the proliferation of *cd44b*^+^ progenitors following SCI and suggested Hapln1-HA-Cd44 signaling is required for stem cell activation and SC repair.

### HA deposition around SC progenitors following SCI requires Hapln1

Hapln1 modulates the levels and stability of HA polymer, we hypothesized that HA is a key effector of Hapln1-HA-Cd44 signaling and that HA-mediated signaling directs progenitor cell proliferation following SCI. We used a biotinylated HA binding protein (HABP) to visualize HA and observed a spectrum of aggregate densities in SC cross sections at 0, 7 and 14 dpi (**Fig. 6A**). In uninjured SCs, HA puncta were the strongest surrounding HuC/D^+^ neurons, consistent with the role of HA in the maintenance of perineural nets. Following SCI, HA aggregates were prominent in the progenitor niche 1 to 2 nuclei radially away from the central canal at 7 dpi. We detected less HA around progenitor cells at 14 dpi, suggesting HA is transiently deposited around SC progenitors in sub-acute SCI. At 7 dpi, HA deposits were significantly decreased in *hapln1a/b* mutants compared to wild-type siblings (**Fig 6B**). These findings showed increased HA deposition around SC progenitors is Hapln1-dependent.

**Figure 6.**
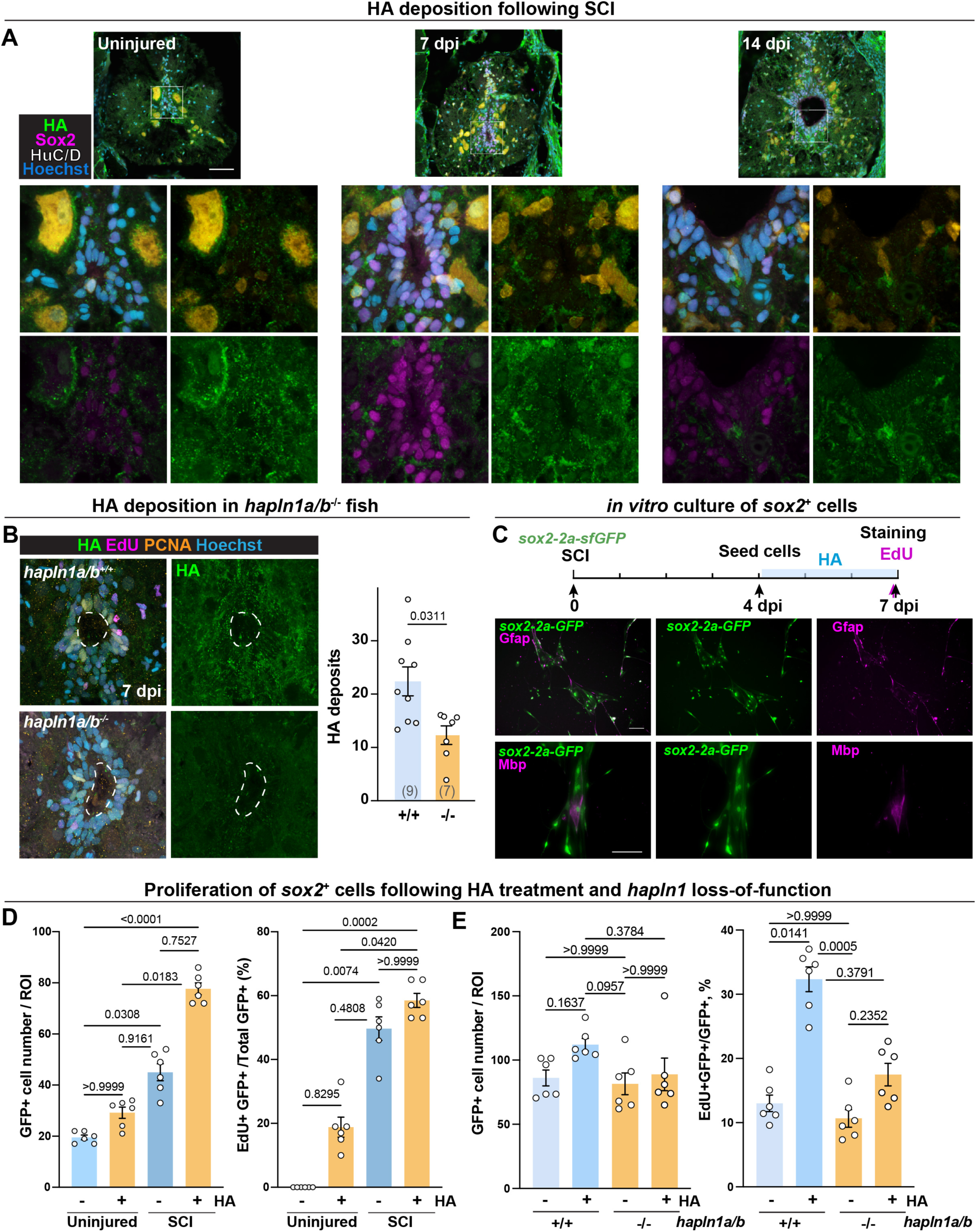
Hapln1 is required for HA deposition and HA-driven progenitor cell proliferation. **(A)** HA binding protein (HABP) staining in wild-type zebrafish at 0, 7 and 14 dpi. Slides were co-stained with Sox2 and HuC/D. **(B)** HAPB staining in *hapln1a/b*^-/-^ and wild-type siblings at 7 dpi. For quantification, integrated density of HABP signal was quantified 450 µm from the lesion. Data points represent individual animals and n numbers are indicated in parentheses. **(C)** *in vitro* culture of primary SC progenitors from zebrafish. SC tissues were dissected from *sox2-2a-sfGFP* at 0 or 4 dpi. Cells were cultured in progenitor media with or without HA. EdU pulse was administered for 4 h before fixation and 3 days post-seeding. The majority of cells were *sox2-2a-sfGFP^+^.* **(D)** Quantification of GFP^+^ cell numbers per ROI (Left) and EdU^+^ GFP^+^ cells relative to GFP^+^ cells (right). Cells collected from 0 or 4 dpi animals were cultured with or without HA. **(E)** Quantification of GFP^+^ cell numbers per ROI (Left) and EdU^+^ GFP^+^ cells relative to GFP^+^ cells (right). Cells collected from *hapln1a/b*^-/-^ or wild-type siblings at 4 dpi animals were cultured with or without HA. Data points indicate 2 biological and 3-4 technical replicates. For SC micrographs, dashed lines delineate the central canal. Scale bars: 50 µm.

### HA promotes Hapln1-dependent proliferation of SC progenitors *in vitro*

As localized HA manipulation to near physiological levels around progenitor cells is technically challenging, we developed an *in vitro* culture model to better understand how Hapln1-HA singaling impacts zebrafish SC progenitors. This method circumvents the constraints of manipulating HA *in vivo* and enables precise manipulation of HA in the culture medium. To culture SC progenitors, we collected SC tissues from adult *sox2-2a-GFP* zebrafish, dissociated and seeded cells in zebrafish CNS progenitor media. Uninjured and 4 dpi SC tissues were collected. Our culture conditions selected for *sox2-2a-GFP*^+^ cells, which accounted for the majority of cells cultured 3 days after plating (**Fig. 6C**). The majority of *sox2-2a-sfGFP*^+^ cells co-stained with Gfap with occasional co-labeling with Mbp (**Fig. 6C**). We therefore used this culture system for HA manipulation.

To examine the effects of HA on progenitor cell proliferation, we cultured SC progenitors and supplemented their cell culture media with 1 mg/mL HA for 3 days (**Fig. 6C**). HA of 1.5 MDa was chosen to better mimic native conditions. Cells were pulsed with EdU for 4 hours prior to fixation and staining. HA treatment induced increased EdU incorporation in *sox2-2a-sfGFP* cells collected from either uninjured or injured tissues (**Fig. 6D**). Importantly, *sox2-2a-sfGFP* cell numbers were only significantly increased in cells cultured following SCI (**Fig. 6D**). We further noted that, in the absence or presence of HA, cells collected from 4 dpi animals were more dense and showed increased EdU incorporation compared to cells collected from uninjured fish (**Fig. 6D**). HA treatment was not sufficient for uninjured cells to match the proliferative profile of cells collected from injured fish. These observations suggested our culture system reflects the proliferative states of the tissues they were collected from, that HA promotes progenitor cell proliferation following SCI, but that HA alone is not sufficient to substitute for the injury signals that prime progenitor cells toward proliferation following injury.

To determine if Hapln1 is required for HA-induced progenitor cell proliferation, we cultured SC progenitors from *hapln1a/b^-/-^; sox2-2a-GFP* or *sox2-2a-GFP* control fish at 4 dpi and assayed their proliferation upon HA supplementation for 3 days. Recapitulating the proliferative effect of HA on SC progenitors, *sox2-2a-GFP^+^* cells from wild-type fish showed a 2.46-fold increase in EdU incorporation. Conversely, EdU incorporation was statistically comparable between *hapln1a/b^-/-^*cells treated with HA compared to *hapln1a/b^-/-^* cells alone (**Fig. 6E**). These results indicated Hapln1 is required for HA-induced progenitor cell proliferation and supported a model in which Hapln1 mediates the effects of HA on progenitor cell proliferation following SCI.

## DISCUSSION

This study uncovers molecular and transcriptional differences between SC progenitors in regenerative zebrafish and non-regenerative mice. Diving into the roles of Hapln1-HA-Cd44 signaling in promoting progenitor cell activation post-injury provided insights into the mechanisms through which zebrafish leverage pro-regenerative ECM components to achieve SC repair.

We propose zebrafish maintain SC progenitors in a stem-like state poised for spontaneous repair, a feature that is not shared with mammalian SC progenitors. Supporting this model, we found zebrafish SC progenitors are more transcriptionally similar to neurogenic progenitors of the mammalian brain rather than non-neurogenic progenitors of the mammalian SC. We also noted that mammalian SC progenitors undergo dramatic molecular changes in response to injury. These changes are consistent with ependymal cells proliferating and differentiating into astrocytes and oligodendrocytes following SCI (Horky et al., 2006; Meletis et al., 2008; Barnabe-Heider et al., 2010; Sabelström et al., 2013). On the other hand, zebrafish SC progenitors can support repair while preserving the gross molecular identities that define them under homeostasic. These findings indicated progenitor cell potency is transcriptionally encoded and suggested SC progenitors are intrinsically pre-equipped with a regenerative transcriptional program. Recognizing the limitations of single-cell sequencing in identifying features that are expressed at low levels or in small number of cells, further cell specific proteomics and *in vivo* validation are needed to fully understand the ECM-mediated mechanisms that support or impair SC regeneration.

ECM has traditionally been associated with scarring and anti-regenerative effects. Extensive data support this model, showing that ECM molecules like Myelin associated glycoproteins (MAGs) and CSPGs are prominent inhibitors of axon regrowth in mammalian SC lesions (Bradbury et al., 2002; Wang et al., 2002; Grimpe and Silver, 2004; Geoffroy and Zheng, 2014). Supporting this model, inhibitory ECM components absent in zebrafish SC lesions impede regeneration when introduced into zebrafish (Kolb et al., 2023). An alternative model posits that inhibitory CSPGs present in zebrafish SCI are leveraged to support repair rather than inhibiting it. Corroborating this model, enzymatic CSPG inhibition improves axon regrowth in mammals but has anti-regenerative effects in zebrafish (Bradbury et al., 2002; Grimpe and Silver, 2004; Kisucká et al., 2026; Lafouasse et al., 2026). Our study supports yet a third model wherein zebrafish progenitors upregulate pro-regenerative ECM-related genes that are absent in mice, highlighting a diversity in the regenerative strategies employed by zebrafish to enact SC repair.

Harnessing fundamental principles from zebrafish SC repair presents new avenues to advance SCI treatments and stem cell strategies. Our findings indicate that, by modulating HA stability, Hapln1 orchestrates Hapln1-HA-Cd44 signaling to enhance progenitor cell responses to injury. Hapln1a and HA deposition are markedly upregulated around zebrafish progenitors following SCI, but not in mouse SC progenitors. These ECM components are known to support neural development and stem cell function (Preston and Sherman, 2011). Intriguingly, HA deposition is prominent around neural stem cells during development but is postnatally decreased, which support its proliferative and neurogenic effects (Margolis et al., 1975). Our data indicate Hapln1 is required for the proliferation of Cd44^+^ progenitors and necessary to mediate HA-driven proliferation *in vitro*. Aligning with its own upregulation in sub-acute SCI, our study establishes Hapln1 as a pivotal player in driving an acute regenerative response post-injury. It remains to be tested whether Hapln1 supplementation is sufficient to reawaken the dormant neurogenic potential of mammalian neural progenitors.

## ACKNOWLEDGMENTS

We thank V. Cavalli and A. Johnson for discussion, J. Wang (Emory University) for zebrafish lines and the Zebrafish Consortium Facility at Washington University for animal care. This research was supported by grants from the NIH (R01 NS113915 and R01 NS123708 to MHM) and the Irving Boime Graduate Fellowship from the Department of Developmental Biology at Washington University School of Medicine (to YX).

## METHODS

### Cross-species integration of single-cell datasets

Mouse datasets used analyzed in this study include a scRNA-seq atlas of mouse SCI (Milich et al., 2021) and a dataset that includes ependymal and progenitor cells from the subventricular zone of uninjured mouse brain (Shah et al., 2018). Zebrafish databases used include a recently generated dataset from sox2-2a-sfGFP fish at 0, 7 and 14 dpi and an atlas of SC regeneration from 0, 7, 21 and 42 dpi (Saraswathy et al., 2024; Weinholtz et al., 2026). For cross-species dataset integration, we first converted zebrafish genes to their mouse orthologues using the Seurat_Species_Converter. This algorithm uses Ensemble biomaRT package for identifying orthologues. The following rules were applied while converting the data matrix: 1) one-to-one orthologue mapping was performed whenever possible, 2) For genes with one to several human orthologues, the corresponding zebrafish RNA data values were copied to every mapped gene in humans, 3) RNA data values were eliminated for zebrafish genes that did not have any human orthologue, 4) RNA data values of paralogous zebrafish genes were added and the cumulative data value was assigned to the human orthologue. After conversion, datasets were integrated and analyzed using Seurat (v4) package with R (v4.3.1) (R core team). Each dataset was independently normalized and scaled using the “SCTransform” function, which is an improved method for normalization, that performs a variance-stabilizing transformation using negative binomial regression. Integration features were selected based on the top 4000 highly variable features using “SelectIntegrationFeatures” function (nfeatures = 4000), which was used as input for the “anchor.features” argument of the “PrepSCTIntegration” function. Integration anchors were identified using the “FindIntegrationAnchors” function with SCT as normalization method. Finally, “IntegrateData” function was used to integrated datasets from each species. PCA analysis was performed on the 4000 variable features and the top 50 principal components selected based on the elbow plot heuristic, which measures the contribution of variation in each component. These 50 principal components were used in “FindNeighbors” and “FindClusters” functions to perform graph-based clustering on a shared nearest neighbor graph(Levine et al., 2015; Xu and Su, 2015). Louvain algorithm was used for modularity optimization in cell clustering using “FindClusters” function. The resolution parameter (res= 0.6 for zebrafish vs mouse SCI, res = 0.9 for zebrafish SC vs mouse brain vSVZ; res = 0.7 zebrafish SC vs mouse SC vs mouse brain vSVZ) that determines the granularity of clustering was selected by visually inspecting clusters with resolutions ranging between 0.1 and 2.0 as well as clustree graphs. Uniform Manifold Approximation and Reduction (UMAP) was used for non-linear dimensional reduction of the first 50 principal components and visualize the data using “RunUMAP” function. Data was graphed using different plot functions, such as “DimPlot”, “VlnPlot”, “FeaturePlot”, “Dotplot”, “plot1cell” and “DoHeatmap”, to view the cell cluster identity and gene expression. Cell number and proportion were extracted using the “table” and “prop.table” functions. Differential gene expression for individual clusters was identified using Wilcoxon rank sum test in the “FindAllMarkers” function. Marker genes detected in at least 25% of the clustered cells and with a logFC threshold of 0.25 were selected. Only positive markers were reported.

### ECM gene expression analysis

To assemble a comprehensive list of ECM-related genes, we compiled the complete list of zebrafish genes from Matrisome.org including 3 core ECM categories (Collagen, ECM glycoproteins and Proteoglycans) and 3 matrisome categories (ECM-affiliated proteins, ECM regulators and Secreted factors). To ensure ECM-related genes were comprehensively included in this analysis, we used Matrisome.org to compile similar lists from mouse and human databases, which were then converted to their zebrafish orthologs using the method described above (Ensemble biomaRT). This analysis generated a list of zebrafish ECM-related genes **(Table S1).** To identify enriched ECM features, single-cell datasets for mouse and zebrafish SCI were separately aggregated using the “AggregateExpression” (Seurat v4.4) function for each time point (Milich et al., 2021; Weinholtz et al., 2026). Differential gene expression analysis was performed for each time point relative to uninjured controls with DESeq2 using a p-value threshold of p<0.05. Genes with |Log2(FC)| > 1 were considered downregulated or upregulated. To compare ECM-related genes between zebrafish and mouse SC progenitors, differentially expressed ECM-related genes were compared and manually verified to match any undetected orthologs **(Table S2).**

### Gene Ontology (GO) analysis

Gene ontology analysis was performed using Metascape. Input and analysis species were set as *D. rerio* for zebrafish dataset and *M. musculus* for mouse and cross-species integrated datasets. Express analysis was performed for gene ontology. Metascape identified statistically enriched terms (including GO biological processes, Reactome gene set and KEGG pathway) and calculated hypergeometric p-values and enrichment factors. Significant terms were hierarchically clustered into a tree based on Kappa-statistical similarities among their gene memberships. A kappa score of 0.3 was applied to cast the tree into term clusters. The most enriched term in each cluster was chosen as the representative term. Dot plots of enriched GO terms were generated in R using “clusterProfiler” package.

### Zebrafish

Adult zebrafish of the Ekkwill/AB strains were maintained at the Washington University Zebrafish Consortium Facility. All animal experiments were performed in compliance with institutional animal protocols. Adult male and female animals of 3-4 cm in length (4-6 months of age) were used. Experimental fish and control siblings of similar size and equal sex distribution were used for all experiments. SC transection surgeries and regeneration analyses were performed in a blinded manner, and at least 2 independent experiments were performed using different clutches of animals. Control and experimental animals in which SC tissues were transected were housed in equal numbers (four to seven fish) in 1.1 L tanks. The following previously published zebrafish strains were used: *sox2-2A-sfGFP^stl84^* (Shin et al., 2014) in addition to *TgBac(hapln1a:eGFP)*, *TgBac(hapln1b:eGFP)*, *TgBac(hapln1a:mCherry-NTR)* and *hapln1a/b^-/-^* (Sun et al., 2022).

### Zebrafish spinal cord injury and treatments

#### Spinal cord transection

Zebrafish were anaesthetized in 0.2 g/l of MS-222 buffered to pH 7.0. Fine scissors were used to make a small incision that transects the spinal cord 4 mm caudal to the brainstem region. Complete transection was visually confirmed at the time of surgery. Injured animals were also assessed at 2 dpi to confirm loss of swim capacity post-surgery.

#### MTZ treatment

To ablate *hapln1a*^+^ cells, *hapln1a*:mCherry-NTR fish and transgene-negative siblings were subjected to complete SC transections, followed by 3 consecutive days of 10 mM MTZ treatment in fish water (Sigma-Aldrich, M1547) for 14 h from 5 to 7 dpi. dsRed and Ccp3 staining were performed at 8 dpi to assess the extents of cell ablation and cell death.

#### EdU labeling

12.5 mM EdU (Sigma 900584) diluted in PBS was used. Zebrafish were anaesthetized as described above, 10 µL of EdU solution was injected intraperitoneally into each zebrafish. Daily injections were performed until experiment end point. Transverse cryosections were generated using method described above. For EdU staining, sections were incubated with freshly prepared EdU staining solution (100 mM Tris at pH 8.5, 1 mM CuSO4, 200 µM Fluorescent azide and 100 mM Ascorbic) for 30 min in the dark at room temperature.

### Swim endurance assays

Zebrafish were exercised in blinded groups of up to 18 fish in a 5 L swim tunnel device (Loligo, SW100605L, 120 V/60 Hz). After 5min of acclimation (with no water flow) inside the enclosed tunnel, fish were subjected to 9 cm/s and 10cm/s of water flow for 5 min each to evaluate their swimming capability at minimum velocity. Water current velocity was then increased every two minutes and fish swam against the current until they reached exhaustion. Exhausted animals were removed from the chamber without disturbing the remaining fish. Swim time at exhaustion was recorded for each fish. Results are expressed as mean ± SEM. For *hapln1a* cell ablation, one-way ANOVA and multiple comparisons were performed using Prism software to determine statistical significance of swim times between groups. For *hapln1a/b* mutants, multiple comparisons from two-way ANOVA were performed.

### Axon regrowth assay

Anterograde axon tracing was performed on adult fish at 14 and 42 dpi. Fish were anesthetized using MS-222 and fine scissors were used to transect the cord 4 mm rostral to the lesion site. Biocytin-soaked 2 mm^3^ gel foam was applied at the new injury site (Gel foam, Pfizer, 09-0315-08; biocytin, saturated solution, Sigma-Aldrich, B4261). Fish were euthanized 3.5 h post-treatment and biocytin was histologically detected using Alexa Fluor 594-conjugated streptavidin (Molecular Probes, S-11227). Biocytin-labeled axons were quantified using the ‘threshold’ and ‘particle analysis’ tools in the Fiji software. Four sections per fish at 600 µm (proximal) and 1500 µm (distal) caudal to the lesion core, and two sections 450 and 750 µm rostral to the lesion, were analyzed. Axon regrowth was calculated by normalizing the efficiency of biocytin labeling rostral to the lesion for each fish. Percentage axon growth was then normalized to that of the control group.

### Glial bridging

GFAP immunohistochemistry was performed on serial transverse sections from adult fish at 14 and 42 dpi. The cross-sectional area of the glial bridge (at the lesion site) and the area of the intact spinal cord (750 µm rostral to the lesion) were measured using FIJI software. Bridging was calculated as a ratio of the bridge area to the rostral area for each fish.

### Histology of zebrafish samples

Transverse cryosections of 16 µm thickness were generated. For immunohistochemistry, tissue sections were rehydrated in PBT, permeabilized with 0.2% TritonX-100 for 5 mi, then blocked with 5% goat serum for 1 h at room temperature. For nuclear antigens such as PCNA and Sox2, antigen retrieval was performed in boiling citrate buffer (pH = 6.0) for 5 min prior to blocking. Sections were incubated in primary antibody solution overnight at 4 °C then washed in PBT and treated for 1 h in secondary antibodies diluted in blocking solution at room temperature. Primary antibodies used in this study were: rabbit anti-Sox2 (GeneTex, 124477, 1:250), mouse anti-Gfap (ZIRC, ZRF1; 1:500), chicken anti-GFP (Aves Labs, GFP-1020; 1:1000), rabbit anti-PCNA (GeneTex, GTX124496; 1:500), mouse anti-HuC/D (Invitrogen, A21271, 1:500), rabbit anti-dsRed (Takara, 632496, 1:250), rabbit anti-Cleaved caspase 3 (Cell Signaling 2810S, 1:250) and Mbp (a kind gift from S. Kucenas, 1:1000). Secondary antibodies used in this study were: Alexa Fluor 488, Alexa Fluor 594, and Alexa 647 goat anti-rabbit or anti-mouse antibodies (Jackson ImmunoResearch, 1:200, 103-545-155, 111-605-003, 111-585-003, 115-605-003 and 115-585-003). Hoechst staining was performed according to the manufacturer’s recommendation (Thermo Fisher Scientific, H3570).

The HCR RNA *in situ* hybridization protocol was adapted from Molecular Instruments. Briefly, tissue sections were hydrated and pre-treated either with 0.2% TritonX-100 for 5 min or boiled in Citrate Buffer (10 mM Citric Acid, 0.05% Tween-20, pH 6.0) for 5 min. For blocking, sections were incubated in Hybridization buffer (Molecular Instruments) for 1 h at 37 °C, then incubated in pre-warmed sets of DNA probes diluted to 0.0015 pmol/uL in Hybridization buffer for 48 h at 37 °C. Washes were then performed with Wash Buffer (Molecular Instruments) at 37 °C followed by 5x SSCT (3 M NaCl, 0.3 M Sodium Citrate, 0.1% Tween-20, pH 7.0). For signal amplification, sections were incubated in Amplification buffer (Molecular Instruments) for 1 h at room temperature. Prior to amplification, h1 and h2 hairpins were snap-cooled separately and diluted 1:50 in Amplification buffer. Amplification proceeded overnight at room temperature in the dark. Samples were then washed in 5x SSCT, 5x SSC and PBT before proceeding to immunohistochemistry. HCR RNA probes used in this study (*hapln1a*, *hapln1b,* and *cd44b*) were designed using a python script and ordered as 50 pmol opools from IDT **(Table S3)**.

For HA staining, a biotinylated HA binding protein (HABP, Sigma #385911, 1:250) was used in conjunction with primary antibody during staining and subsequently visualized by Alexa Fluor 488-conjugated streptavidin (Molecular Probes, S11223).

Tissue sections were imaged using a Zeiss LSM 800 confocal microscope or a Zeiss Axioscan.Z1 slide scanner for HCR *in situ* hybridization and immunofluorescence.

### Quantification

#### HCR in situ hybridization quantification

Images were converted into a 16-bit tiff; the spinal cord cross sections were outlined as region of interest (ROI); comparable thresholds were set and recorded for each image; and areas past the threshold were measured. Area (px^2), % of area relative to the whole spinal cord area, and integrated density of the HCR signal were recorded. Ordinary two-way ANOVA were performed using Prism software to determine statistical significance between timepoints.

#### hapln1a and sox2-2a-sfGFP colocalization quantification

HCR *in situ* hybridization signals were quantified using a customized Fiji script (Saraswathy et al., 2024). SC perimeters were outlined as ROI. Thresholds were user-defined and recorded for the *hapln1a* channel and *sox2-2a-sfGFP* channel to set to create an inverted mask. Integrated density was quantified using the “Analyze Particles” command. To quantify RNA signals inside *sox2-2a-sfGFP*^+^ cells, an inverted mask was created after thresholding *sox2-2a-sfGFP* ^+^ cells, then added to the final quantification mask. Fluorescence signals inside this newly generated mask were considered *sox2-2a-sfGFP* ^+^ cell specific. We measured the ratio of the summed intensities of pixels from the *sox2-2a-sfGFP* channel for which the intensity in the *hapln1a* channel is above zero to the total intensity in the *hapln1a* channel.

#### Cell counting

Counting of EdU^+^, Sox2^+^, PCNA^+^ and *cd44b+* cells were performed using a customized Fiji script (adapting ITCN: Image based Tool for counting nuclei; https://imagej.nih.gov/ij/plugins/itcn.html). The Fiji script incorporated user-defined inputs to define channels (including Hoechst) and to outline ROI. To quantify nuclei, the following parameters were set in ITCN counter: width, 15; minimal distance, 7.5; threshold, 0.3. Once nuclei were identified, user-defined thresholds of individual cell markers were used to mask the image and identify nuclei located inside the masked regions. *xy* coordinates were extracted for each nucleus for cell counting. Raw counts and *xy* coordinates from Fiji were processed using a customized R script. Colocalization analyses were performed using the JACoP plugin in Fiji. Markers were considered overlapping if they shared nuclei with same *xy* coordinates.

For all statistical data, data points represent individual animals and N numbers are indicated. Statistical tests are reported in figure legends for each panel. Multiple comparisons were performed within groups and significant observations were reported.

### Cell culture

#### Zebrafish spinal cord progenitor culture

Zebrafish SC tissues were dissected and dissociated in 0.25% Trypsin EDTA (0.25%) (Invitrogen cat# 25200056). For each 35 mm dish, 4 x 10^5^ cells were seeded and left unagitated for 16-24h post-seeding. Complete progenitor medium consists of Lebovitz-15 medium (ThermoFisher cat# 11415064) supplemented with 5% fetal bovine serum (Sigma #F2442), 1x L-Glutamine (Invitrogen #25030081), 1% N2, 1% B27 with vitamin A, and 500 U/mL of penicillin-streptomycin (ThermoFisher cat# 15070063). 50% of the culture medium was changed every day until sample fixation and immunofluorenscence. For HA supplementation, high molecular weight hyaluronic acid (Sigma #**53747)** was first UV-sterilized and reconstituted in complete progenitor medium.

#### Human iPSC-derived neuroepithelial (NEP) cell culture

The human iPSC line AN1.1 used in this study was obtained from the Human Cells, Tissues, and Organoids Core (hCTO) at Washington University in St. Louis. The AN1.1 line was derived from renal epithelial cells of a healthy Asian female donor through reprogramming at Washington University in St. Louis. iPSCs were maintained on Matrigel-coated plates in mTeSR Plus medium (STEMCELL, #100-0276). To generate NEPs, iPSCs were dissociated with 1x TrypLE (ThermoFisher Scientific, #15140122) and plated at 1.5 x 10^5^ cells/well in Matrigel-coated plates. Cells were cultured in a base medium consisting of 1:1 DMEM/F12 (Gibco) and Neurobasal medium(Gibco), 0.5x N2 Supplement (Gibco), 0.5x B27 Plus Supplement (Gibco), 0.1 mM ascorbic acid (Santa Cruz), 1x GlutaMax (Gibco) and 1x penicillin/streptomycin (Gibco) supplemented with 3 μM CHIR99021 (STEMCELL), 2 μM DMH1 (STEMCELL), 2 μM SB431542 (Fisher Scientific), and 10 μM ROCK inhibitor (STEMCELL). On the following day, the medium was replaced with NEP induction medium without ROCK inhibitor for maintenance. For HA supplementation experiment, cells were seeded at low density and cultured for 4 days before fixation.

#### Staining and imaging

Immunofluorescence and EdU staining were performed as described above. Images were acquired using the EVOS M7000 Imaging system (ThermoFisher). Besides image analyses methods described above, zebrafish progenitor culture staining were manually quantified using the FIJI built-in “Cell Counter.” Manually marked cells were recorded for each ROI. For each experiment, at least 2 biological replicates and 3 technical replicate images from different ROIs were analyzed. For all statistical data, data points represent individual values. Statistical tests are reported in figure legends for each panel. Multiple comparisons were performed within groups and significant observations were reported.

## REFERENCES

Barnabe-Heider, F., Goritz, C., Sabelstrom, H., Takebayashi, H., Pfrieger, F. W., Meletis, K. and Frisen, J. (2010) ‘Origin of new glial cells in intact and injured adult spinal cord’, Cell Stem Cell 7(4): 470–82.

Becker, C. G. and Becker, T. (2008) ‘Adult zebrafish as a model for successful central nervous system regeneration’, Restorative neurology and neuroscience 26(2-3): 71–80.

Bradbury, E. J., Moon, L. D., Popat, R. J., King, V. R., Bennett, G. S., Patel, P. N., Fawcett, J. W. and McMahon, S. B. (2002) ‘Chondroitinase ABC promotes functional recovery after spinal cord injury’, Nature 416(6881): 636–40.

Burris, B., Jensen, N. and Mokalled, M. H. (2021) ‘Assessment of Swim Endurance and Swim Behavior in Adult Zebrafish’, Journal of visualized experiments: JoVE(177).

Burris, B. and Mokalled, M. H. (2024) ‘Spinal Cord Injury and Assays for Regeneration’, Methods Mol Biol 2707: 215–222.

Curado, S., Stainier, D. Y. and Anderson, R. M. (2008) ‘Nitroreductase-mediated cell/tissue ablation in zebrafish: a spatially and temporally controlled ablation method with applications in developmental and regeneration studies’, Nat Protoc 3(6): 948–54.

Cyphert, Jaime M., Trempus, Carol S. and Garantziotis, Stavros (2015) ‘Size Matters: Molecular Weight Specificity of Hyaluronan Effects in Cell Biology’, International Journal of Cell Biology 2015: 563818–563818.

Geoffroy, C. G. and Zheng, B. (2014) ‘Myelin-associated inhibitors in axonal growth after CNS injury’, Current opinion in neurobiology 27: 31–8.

Grimpe, B. and Silver, J. (2004) ‘A novel DNA enzyme reduces glycosaminoglycan chains in the glial scar and allows microtransplanted dorsal root ganglia axons to regenerate beyond lesions in the spinal cord’, J Neurosci 24(6): 1393–7.

Grycz, K., Glowacka, A., Ji, B., Krzywdzinska, K., Charzynska, A., Czarkowska-Bauch, J., Gajewska-Wozniak, O. and Skup, M. (2022) ‘Regulation of perineuronal net components in the synaptic bouton vicinity on lumbar alpha-motoneurons in the rat after spinalization and locomotor training: New insights from spatio-temporal changes in gene, protein expression and WFA labeling’, Exp Neurol 354: 114098.

Horky, L. L., Galimi, F., Gage, F. H. and Horner, P. J. (2006) ‘Fate of endogenous stem/progenitor cells following spinal cord injury’, J Comp Neurol 498(4): 525–38.

Hui, S. P., Dutta, A. and Ghosh, S. (2010) ‘Cellular response after crush injury in adult zebrafish spinal cord’, Dev Dyn 239(11): 2962–79.

Jones, Leonard L., Margolis, Richard U. and Tuszynski, Mark H. (2003) ‘The chondroitin sulfate proteoglycans neurocan, brevican, phosphacan, and versican are differentially regulated following spinal cord injury’, Experimental Neurology 182(2): 399–411.

Kisucká, Alexandra, Bimbová, Katarína Kiss, Kuruc, Tomáš, Ileninová, Mária, Kuchárová, Karolína, Ihnátová, Lenka, Bačová, Mária, Magurová, Martina, Gálik, Ján and Lukáčová, Nadežda (2026) ‘Molecular Changes in CSPG and Glial Scar Markers in Response to Subpial Chondroitinase ABC Treatment Following Spinal Cord Injury’, European Journal of Neuroscience 63(1): e70389–e70389.

Klatt Shaw, D., Saraswathy, V. M., Zhou, L., McAdow, A. R., Burris, B., Butka, E., Morris, S. A., Dietmann, S. and Mokalled, M. H. (2021) ‘Localized EMT reprograms glial progenitors to promote spinal cord repair’, Developmental cell 56(5): 613–626 e7.

Kolb, J., Tsata, V., John, N., Kim, K., Mockel, C., Rosso, G., Kurbel, V., Parmar, A., Sharma, G., Karandasheva, K. et al. (2023) ‘Small leucine-rich proteoglycans inhibit CNS regeneration by modifying the structural and mechanical properties of the lesion environment’, Nat Commun 14(1): 6814.

Lafouasse, L., Koutsogiannis, K., Dai, Y. E., Del Vecchio, L., Pedroni, A., Tsagkogiannis, D., Habicher, J. and Ampatzis, K. (2026) ‘Time-dependent adaptations of damaged neurons and their microenvironment in the regenerating adult zebrafish spinal cord’, Sci Adv 12(10): eaea2882.

Lemons, Michele L., Howland, Dena R. and Anderson, Douglas K. (1999) ‘Chondroitin Sulfate Proteoglycan Immunoreactivity Increases Following Spinal Cord Injury and Transplantation’, Experimental Neurology 160(1): 51–65.

Liu, H., Li, T., Ma, B., Wang, Y. and Sun, J. (2023) ‘Hyaluronan and Proteoglycan Link Protein 1 Activates the BMP4/Smad1/5/8 Signaling Pathway to Promote Osteogenic Differentiation: an Implication in Fracture Healing’, Mol Biotechnol.

Long, Katherine R., Newland, Ben, Florio, Marta, Kalebic, Nereo, Langen, Barbara, Kolterer, Anna, Wimberger, Pauline and Huttner, Wieland B. (2018) ‘Extracellular Matrix Components HAPLN1, Lumican, and Collagen I Cause Hyaluronic Acid-Dependent Folding of the Developing Human Neocortex’, Neuron 99(4): 702–719.e7.

Mahony, C. B., Cacialli, P., Pasche, C., Monteiro, R., Savvides, S. N. and Bertrand, J. Y. (2021) ‘Hapln1b, a central organizer of the ECM, modulates kit signaling to control developmental hematopoiesis in zebrafish’, Blood Adv 5(23): 4935–4948.

Margolis, R. U., Margolis, R. K., Chang, L. B. and Preti, C. (1975) ‘Glycosaminoglycans of brain during development’, Biochemistry 14(1): 85–8.

Meletis, K., Barnabe-Heider, F., Carlen, M., Evergren, E., Tomilin, N., Shupliakov, O. and Frisen, J. (2008) ‘Spinal cord injury reveals multilineage differentiation of ependymal cells’, PLoS Biol 6(7): e182.

Milich, L. M., Choi, J. S., Ryan, C., Cerqueira, S. R., Benavides, S., Yahn, S. L., Tsoulfas, P. and Lee, J. K. (2021) ‘Single-cell analysis of the cellular heterogeneity and interactions in the injured mouse spinal cord’, J Exp Med 218(8).

Mokalled, M. H., Patra, C., Dickson, A. L., Endo, T., Stainier, D. Y. and Poss, K. D. (2016) ‘Injury-induced ctgfa directs glial bridging and spinal cord regeneration in zebrafish’, Science 354(6312): 630–634.

Naba, Alexandra, Clauser, Karl R., Hoersch, Sebastian, Liu, Hui, Carr, Steven A. and Hynes, Richard O. (2012) ‘The matrisome: In silico definition and in vivo characterization by proteomics of normal and tumor extracellular matrices’, Molecular and Cellular Proteomics 11(4).

Nauroy, Pauline, Hughes, Sandrine, Naba, Alexandra and Ruggiero, Florence (2018) ‘The in-silico zebrafish matrisome: A new tool to study extracellular matrix gene and protein functions’, Matrix Biology 65: 5–13.

Preston, M. and Sherman, L. S. (2011) ‘Neural stem cell niches: roles for the hyaluronan-based extracellular matrix’, Front Biosci (Schol Ed) 3(3): 1165–79.

Reimer, M. M., Sorensen, I., Kuscha, V., Frank, R. E., Liu, C., Becker, C. G. and Becker, T. (2008) ‘Motor neuron regeneration in adult zebrafish’, The Journal of neuroscience: the official journal of the Society for Neuroscience 28(34): 8510–6.

Sabelström, Hanna, Stenudd, Moa, Réu, Pedro, Dias, David O., Elfineh, Marta, Zdunek, Sofia, Damberg, Peter, Göritz, Christian and Frisén, Jonas (2013) ’Resident Neural Stem Cells Restrict Tissue Damage and Neuronal Loss After Spinal Cord Injury in Mice’, *Science (New York*, N.Y.) 342(6158): 637–640.

Sanchez-Ventura, J., Lane, M. A. and Udina, E. (2022) ‘The Role and Modulation of Spinal Perineuronal Nets in the Healthy and Injured Spinal Cord’, Frontiers in cellular neuroscience 16: 893857.

Saraswathy, V. M., Zhou, L., McAdow, A. R., Burris, B., Dogra, D., Reischauer, S. and Mokalled, M. H. (2022) ‘Myostatin is a negative regulator of adult neurogenesis after spinal cord injury in zebrafish’, Cell Rep 41(8): 111705.

Saraswathy, V. M., Zhou, L. and Mokalled, M. H. (2024) ‘Single-cell analysis of innate spinal cord regeneration identifies intersecting modes of neuronal repair’, Nat Commun 15(1): 6808.

Shah, P. T., Stratton, J. A., Stykel, M. G., Abbasi, S., Sharma, S., Mayr, K. A., Koblinger, K., Whelan, P. J. and Biernaskie, J. (2018) ‘Single-Cell Transcriptomics and Fate Mapping of Ependymal Cells Reveals an Absence of Neural Stem Cell Function’, Cell 173(4): 1045–1057 e9.

Shin, J., Chen, J. and Solnica-Krezel, L. (2014) ‘Efficient homologous recombination-mediated genome engineering in zebrafish using TALE nucleases’, Development 141(19): 3807–18.

Silver, Jerry (2016) ‘The glial scar is more than just astrocytes’, Experimental Neurology 286: 147–149.

Sofroniew, Michael V. (2018) ‘Dissecting spinal cord regeneration’, Nature 557(7705): 343–350.

Sun, J., Peterson, E. A., Chen, X. and Wang, J. (2023) ‘hapln1a(+) cells guide coronary growth during heart morphogenesis and regeneration’, Nat Commun 14(1): 3505.

Sun, J., Peterson, E. A., Wang, A. Z., Ou, J., Smith, K. E., Poss, K. D. and Wang, J. (2022) ‘hapln1 Defines an Epicardial Cell Subpopulation Required for Cardiomyocyte Expansion During Heart Morphogenesis and Regeneration’, Circulation 146(1): 48–63.

Tendolkar, A. and Mokalled, M. H. (2025) ‘Mechanisms underpinning spontaneous spinal cord regeneration’, Development 152(20).

Wang, K. C., Koprivica, V., Kim, J. A., Sivasankaran, R., Guo, Y., Neve, R. L. and He, Z. (2002) ‘Oligodendrocyte-myelin glycoprotein is a Nogo receptor ligand that inhibits neurite outgrowth’, Nature 417(6892): 941–4.

Weinholtz, Chase A., Zhou, Lili, Saraswathy, Vishnu Muraleedharan, Xu, Yuxiao, Shaw, Dana Klatt, McAdow, Anthony R., Park, Dongkook, Shin, Jimann, Solnica-Krezel, Lila, Johnson, Aaron N et al. (2026) ‘Transient activation of potent progenitor cells is required for spinal cord regeneration’, bioRxiv: 2026.02.04.703854.

Wolf, K. J., Shukla, P., Springer, K., Lee, S., Coombes, J. D., Choy, C. J., Kenny, S. J., Xu, K. and Kumar, S. (2020) ‘A mode of cell adhesion and migration facilitated by CD44-dependent microtentacles’, Proc Natl Acad Sci U S A 117(21): 11432–11443.

Zheng, B. and Tuszynski, M. H. (2023) ‘Regulation of axonal regeneration after mammalian spinal cord injury’, Nat Rev Mol Cell Biol 24(6): 396–413.

Zhou, L., McAdow, A. R., Yamada, H., Burris, B., Klatt Shaw, D., Oonk, K., Poss, K. D. and Mokalled, M. H. (2023) ‘Progenitor-derived glia are required for spinal cord regeneration in zebrafish’, Development 150(10).

