## Supplemental Figures for "Hapln1-HA signaling promotes progenitor cell proliferation and spinal cord regeneration"

### Figure S1

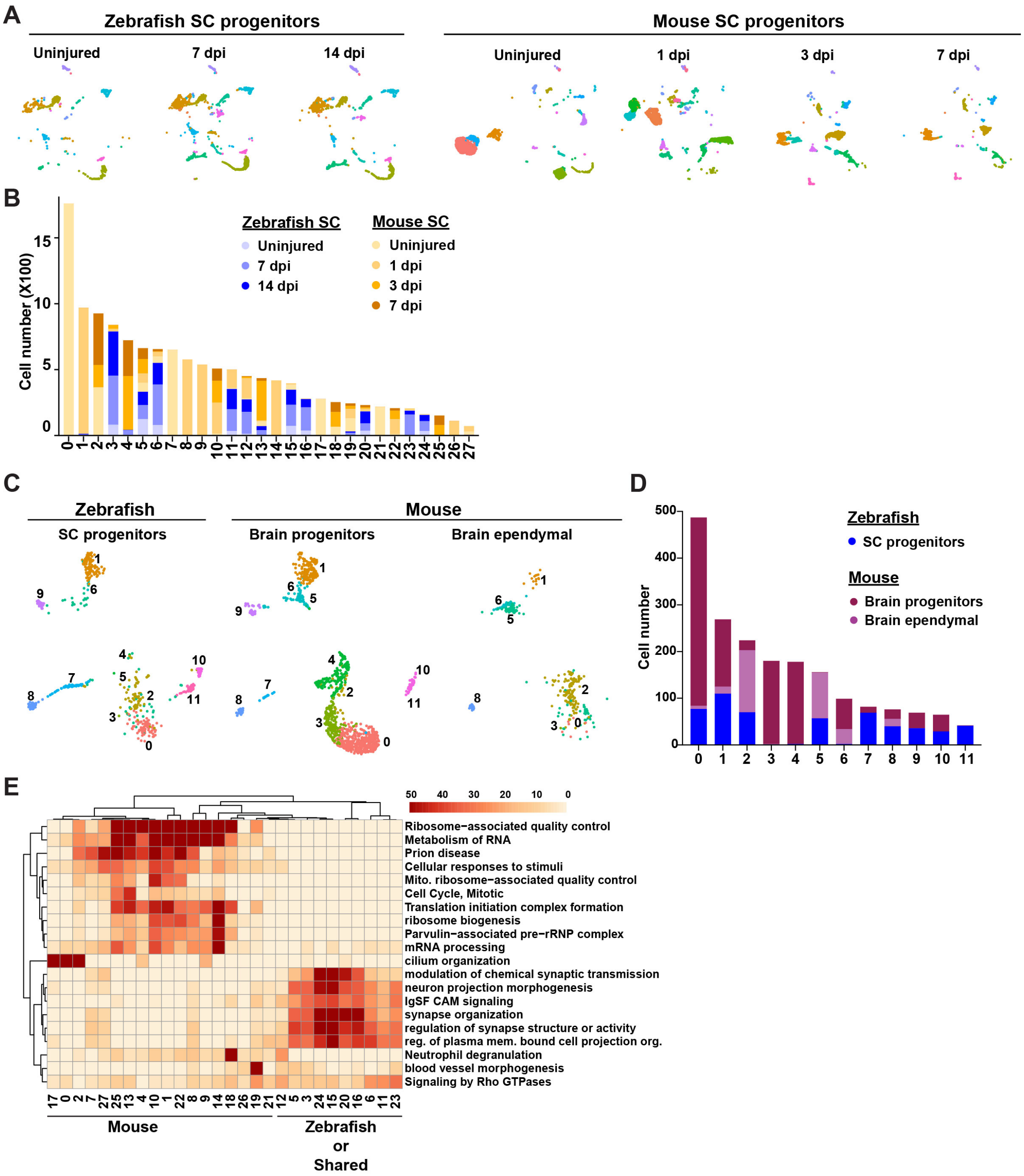


**Figure S1. Comparative transcriptomics between zebrafish and mouse SC progenitors. (A)** Split UMAP of the integrated dataset of *sox2*⁺ cells from zebrafish and mouse SCI. **(B)** Numbers of cells within each cluster from the integrated dataset of *sox2*⁺ cells from zebrafish and mouse SCI. **(C)** Split UMAP of the integrated dataset of *sox2*⁺ cells from uninjured zebrafish SCs with progenitor and ependymal cells from mouse brain. **(D)** Numbers of cells within each cluster from the integrated dataset of *sox2*⁺ cells from uninjured zebrafish SCs with progenitor and ependymal cells from mouse brain. **(E)** Clustered dendrogram and heatmap representation GO terms enriched in each cluster. Top markers from each cluster were used as input.

#### Figure S2

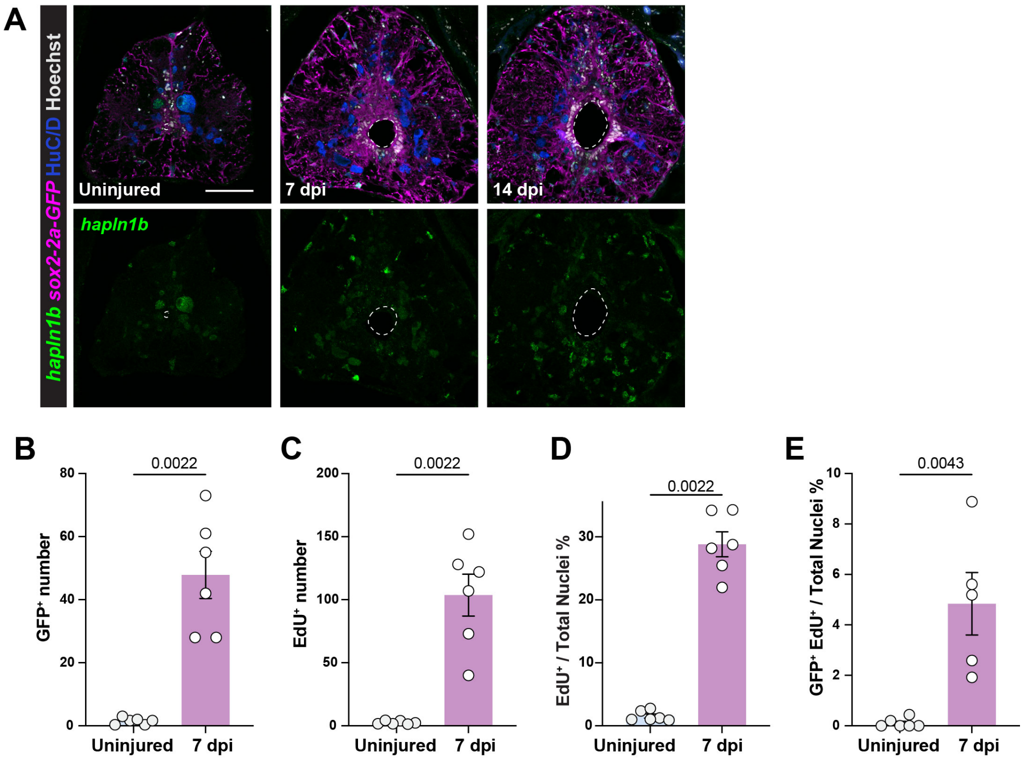


**Figure S2. (A)** HCR *in situ* hybridization for *hapln1b in sox2-2a-sfGFP* fish. SC cross sections at 0, 7 and 14 dpi are shown and co-stained with HuC/D. **(B-E)** Cell proliferation in *hapln1a:GFP* fish following SCI. Quantification of $GFP^+$ cell numbers (B), $EdU^+$ cell numbers and proportions (C,D), and $GFP^+$ $EdU^+$ cell proportions (E) at 0 and 7 dpi.

#### Figure S3

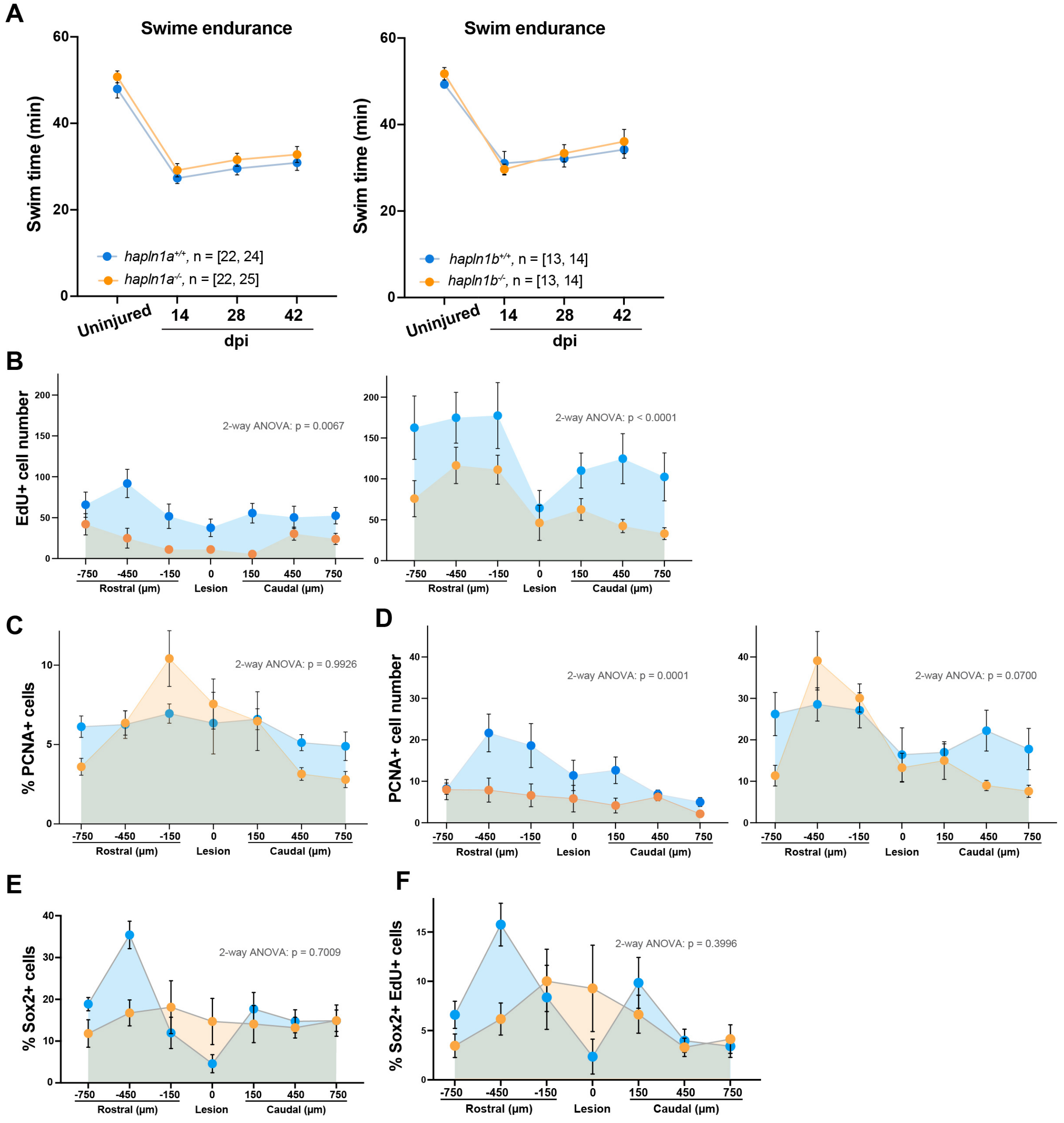


**Figure S3. (A)** Swim endurance in *hapln1a* $^{-/-}$ (Left) or *hapln1b* $^{-/-}$ (right) fish compared to their respective wild-type siblings. **(B-F)** EdU incorporation in *hapln1a/b* mutants. Quantification of EdU incorporation (B), PCNA$^{+}$ cells (C,D), Sox2$^{+}$ cells (E) and Sox2$^{+}$ EdU$^{+}$ cells (F) at 7 or 14 dpi. Multiple tissue levels 750 µm rostral and caudal to the lesion were quantified. Statistical tests and p-values are indicated.

### Figure S4

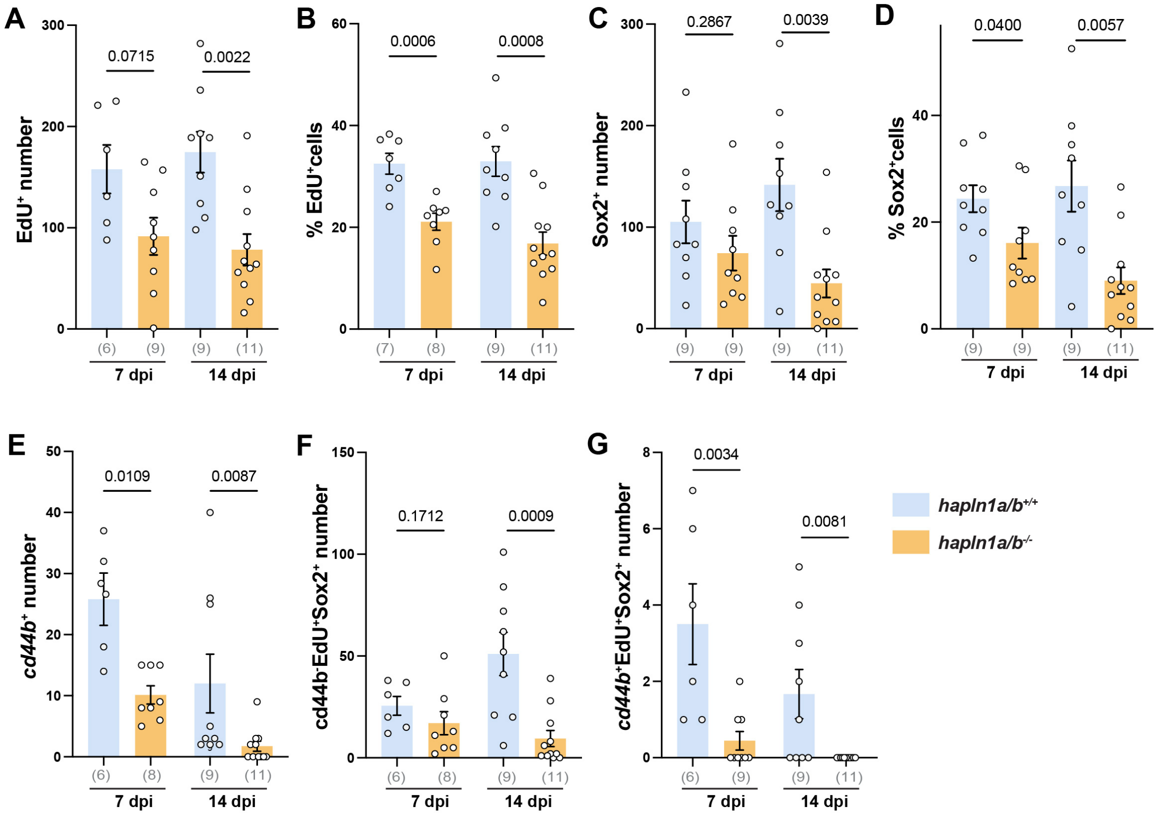


**Figure S4. (A-G)** Proliferation of *cd44b*$^{+}$ cells in *hapln1a/b*$^{-/-}$ and wild-type siblings at 7 dpi. EdU incorporation (A,B), Sox2$^{+}$ cells (C,D), *cd44b*$^{+}$ cells (E), *cd44b*$^{-}$ EdU$^{+}$ Sox2$^{+}$ cells (F) and *cd44b*$^{+}$ EdU$^{+}$ Sox2$^{+}$ cells (G) were quantified. Data points represent individual animals, n numbers are indicated in parentheses, and p-values are indicated.
